# Methionine Deprivation-induced Reprogramming of Hepatic Rhythms Is Mediated by Glucocorticoid Receptor

**DOI:** 10.64898/2026.05.04.722637

**Authors:** Yang Liu, Sara A. Grimm, Fred B. Lih, Leesa J. Deterding, Robert H. Oakley, John A. Cidlowski, Paul A. Wade

**Affiliations:** Eukaryotic Transcriptional Regulation Group, Epigenetics & RNA Biology Laboratory, National Institute of Environmental Health Sciences, Research Triangle Park, NC 27709, USA; Integrative Bioinformatics Support Group, Biostatistics and Computational Biology Laboratory, National Institute of Environmental Health Sciences, Research Triangle Park, NC 27709, USA; Mass Spectrometry Research & Support Group, Epigenetics & RNA Biology Laboratory, National Institute of Environmental Health Sciences, Research Triangle Park, NC, 27709, USA; Molecular Endocrinology Group, Molecular & Cellular Biology Laboratory, National Institute of Environmental Health Sciences, Research Triangle Park, NC 27709, USA

**Keywords:** Circadian rhythms, Methionine starvation, Chromatin, Glucocorticoid receptor, Lipid metabolism

## Abstract

The circadian clock tightly interacts with cellular metabolism, and nutritional challenges can profoundly remodel its function. Dietary methionine restriction has been shown to improve metabolic health and extend lifespan. However, whether and how dietary methionine restriction impacts the circadian clock is unclear. Here, we demonstrate that methionine deprivation (MD) in the mouse induces alterations in multiple signaling systems, inducing rhythmic biological events in liver that are entrained by feeding time rather than day/light cycles. Alteration in growth factor signaling leads to loss of activation of a pioneer transcription factor, STAT5, which normally directs rhythmic chromatin localization of the glucocorticoid receptor (GR). Loss of active STAT5 leads to reprogramming of the epigenome and of the GR-directed transcriptome in a manner abolished by liver-specific deletion of GR. These results demonstrate that dietary methionine restriction remodels the circadian biological program directed by glucocorticoid receptor.

## INTRODUCTION

Circadian rhythms play a crucial role in regulating various physiological processes, including the sleep-wake cycle, glucose metabolism, and energy homeostasis^1^. Disruptions in circadian alignment due to genetic and environmental factors can increase the risk of metabolic diseases such as hepatic steatosis, diabetes, obesity, dyslipidemia, and hyperglycemia^2,3^. Conversely, cellular metabolism is essential for maintaining the circadian clock. Emerging evidence indicates that certain metabolites directly or indirectly regulate circadian oscillators by influencing epigenetic modifications, protein modifications, protein turnover, or protein-DNA interactions^4–7^. Moreover, numerous studies have demonstrated that modifications in dietary composition, such as ketogenic diets and high-fat diets (HFD), can significantly impact peripheral clocks^8–10^.

Methionine metabolism has a profound effect on the methylation status of proteins, nucleic acids, and metabolites, as the intermediate metabolite S-adenosyl methionine (SAM) serves as the primary methyl donor during transmethylation processes^11,12^. Methionine restriction leads to altered methylation by reducing SAM concentration. Methionine starvation activates histone demethylation through the hyperphosphorylation of specific demethylase enzymes, which is regulated by the demethylation of the protein phosphatase PP2A^13^. Recent studies have revealed that metabolites, both methionine and SAM, in the methionine metabolism pathway exhibit circadian oscillations in mouse liver^14^. Moreover, methylation of DNA, RNA, and histones has been reported to regulate the circadian clock, affecting transcription and RNA processing^15–17^. These observations underscore the complex interplay between methionine metabolism and circadian networks.

Chronic dietary methionine restriction with a 75-80% reduction in methionine has been shown to extend lifespan and improve metabolic health in rodents^18,19^. Dietary methionine restriction leads to increased serum levels of fibroblast growth factor 21 (FGF21), which acts as a critical mediator to regulate energy balance^20,21^. Methionine restriction also exerts its influence on numerous biological processes through alterations in histone and RNA methylation^22–25^. Interestingly, dietary methionine restriction has been reported to restore disrupted diurnal metabolism in obese mice fed a high-fat diet^26^. However, our understanding of how dietary methionine influences the circadian clock and the underlying mechanisms remains limited. Recent studies have shown that complete methionine deprivation (MD) regimens can effectively and rapidly promote metabolic health, accompanied by significant alterations in histone modifications in the liver^27,28^.

Here, we show that short-term dietary methionine starvation induces a significant reprogramming of circadian rhythms. MD diet reshapes animal physiology, including oscillation of circulating hormones, such as FGF21 and corticosterone. Cyclic transcripts in liver were rewired by MD diet. Notably, the oscillation of sterol regulatory element-binding transcription factor 1 (SREBF1), a master regulator of de novo lipogenesis, is compromised by MD diet. We also observed MD diet-induced de novo oscillation of thousands of transcripts, entrained by the feeding cycle and accompanied by reshaping of oscillating histone modifications in liver. Finally, we reveal that MD diet abolishes the normal circadian oscillation in signal transducer and activator of transcription 5 (STAT5) signaling, remodels the rhythmic GR binding to chromatin, and alters rhythmic H3K27ac occupancy. These studies provide insights into the molecular mechanisms underlying the nutritional regulation of circadian rhythms.

## RESULTS

### Short-term MD diet resets circadian metabolism

The metabolites and metabolic enzymes in the methionine metabolism pathway, which are closely associated with the regulation of circadian clock^29^, are regulated by dietary methionine availability (Figure 1A). To investigate the impact of dietary methionine on circadian rhythms, six-month-old C57/B6J male mice were assigned to either a control amino acid diet containing 0.82% methionine or a complete methionine deprivation (MD) diet containing 0% methionine^28^. Mice consuming the MD diet exhibited a rapid and significant loss of body weight (Figure S1A). The reduced body lean and fat mass were observed in mice on MD diet for two weeks (Figure S1B). Notably, after one week on the MD diet, mice exhibited a significant reduction in fasting glucose levels and improved glucose tolerance and insulin sensitivity (Figure S1C). Liver function tests were performed on mice fed the MD diet for three weeks. While there were no changes in alkaline phosphatase (ALP) and sorbitol dehydrogenase (SDH) levels between mice on the control diet and MD diet, mice fed the MD diet showed higher levels of alanine aminotransferase (ALT) and aspartate aminotransferase (Figure S1D). Both serum HDL (high-density lipoprotein) and LDL (low-density lipoprotein) cholesterol levels were reduced by the MD diet, whereas serum bile acid levels were elevated in MD diet-fed mice (Figures S1E and S1F). These findings suggest that short-term MD diet affects the metabolic functions of liver.

**Figure 1.**
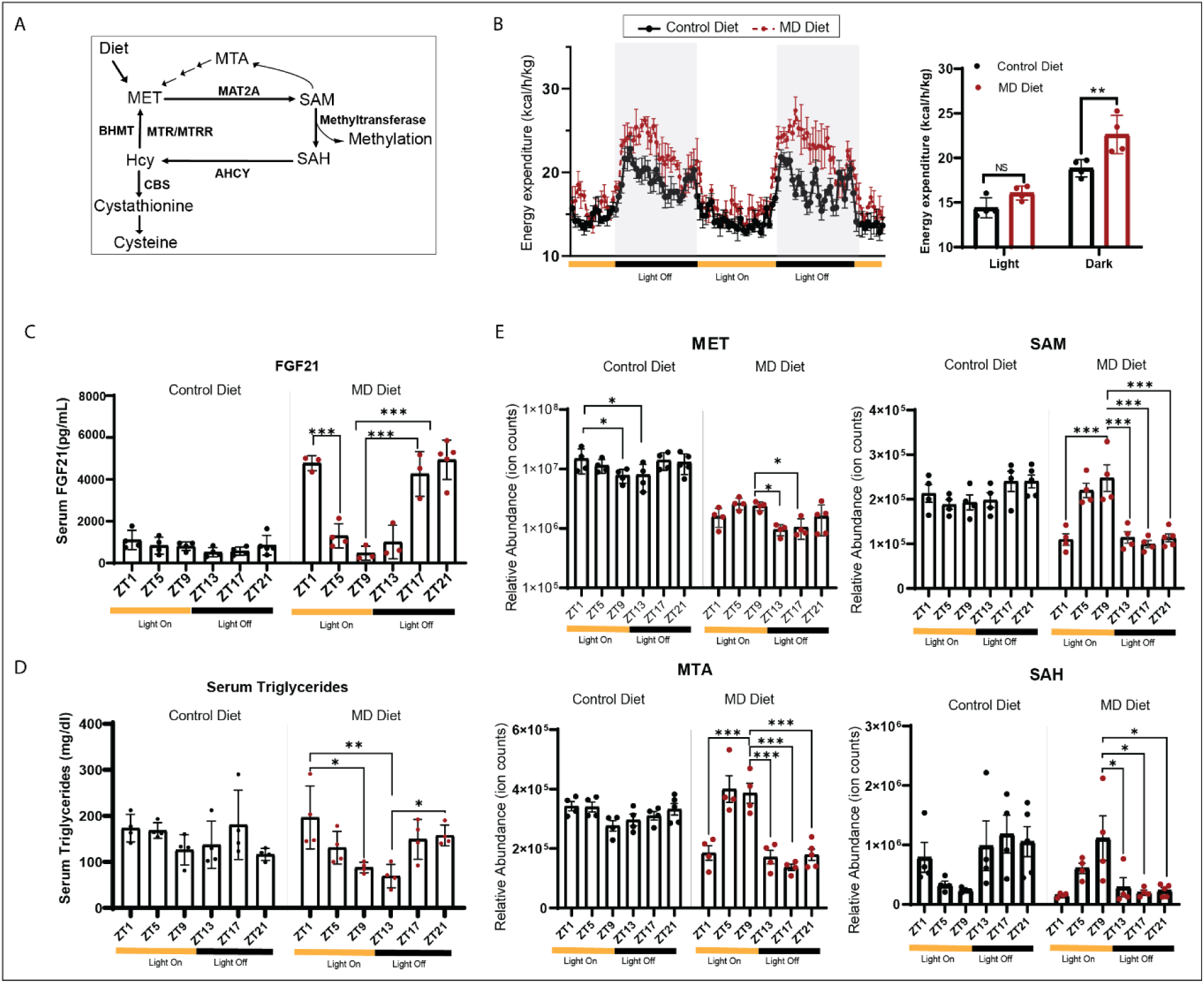
Short-term Methionine deprivation reshapes circadian metabolism. A. The chart depicts key enzymes and metabolites in the methionine cycle. B. On the left, energy expenditure, normalized to body mass, of six-month old male mice after 1 week on the indicated diet. The light/dark cycle in the facility is indicated by shading and on the axis. On the right, quantitation of normalized energy expenditure during dark and light periods is indicated in the column graph (4 animals per group). Significance (**p<0.01, two-way ANOVA with Tukey’s multiple comparisons test) is indicated with double asterisks. C. Serum levels of FGF21 of six-month old male mice after three weeks on the indicated diet. The column graphs indicate mean value for 3-5 animals at indicated zeitgeber time points. Significance (*p<0.05, ***p<0.001, one-way ANOVA with Tukey’s multiple comparisons test) is indicated with asterisks. D. Serum TG of six-month old male mice after three weeks on the indicated diet. The column graphs indicate mean values for 4 animals at indicated zeitgeber time points. Significance (*p<0.05, **p<0.01, one-way ANOVA with Tukey’s multiple comparisons test) is indicated with asterisks. E. Serum levels of Methionine, SAM, MTA, and SAH of six-month old male mice after three weeks on the indicated diet. The column graphs indicate mean values for 4-5 animals at indicated zeitgeber time points. Significance (*p<0.05, ***p<0.001, one-way ANOVA with Tukey’s multiple comparisons test) is indicated with asterisks. See also Figure S1.

To explore the effects of short-term methionine starvation on energy balance, we measured the energy expenditure of mice after one week on the diet. The results demonstrated that mice fed the MD diet exhibited higher energy expenditure compared to mice on the control diet. Notably, this increase in energy expenditure was specifically observed during the dark phase, the active period for mice (Figure 1B), indicating that the MD diet may influence diurnal energy metabolism.

The serum from mice after three weeks on the control and MD diet was collected every four hours throughout a 24-hour light-dark cycle for measuring the levels of circulating hormones and metabolites. We noted that the serum FGF21 levels exhibited an oscillatory pattern in mice fed MD diet, whereas there was no oscillation of FGF21 levels observed in mice fed on control diet (Figure 1C). The elevated circulating FGF21 levels were specifically detected at zeitgeber time (ZT)17, ZT21 and ZT1 in mice fed on MD diet. However, its levels in mice on the MD diet were comparable to those of mice on the control diet at ZT5, ZT9 and ZT13 (Figure 1C). Interestingly, we also observed oscillatory serum triglyceride levels specifically in mice on the MD diet (Figure 1D). The levels of four metabolites, including methionine, SAM, 5-methylthioadenosine (MTA), and S-adenosyl homocysteine (SAH) in methionine metabolism were measured. In comparison to mice on the control diet, mice fed the MD diet experienced a significant decline in serum methionine levels, with a reduction ranging from 70-90%. Interestingly, we noticed that mice on the MD diet exhibited a pronounced reduction in methionine levels specifically during the dark phase, showing a significant difference between ZT13 and ZT9 (Figure 1E). Moreover, we observed circadian oscillation of SAM in mice on the MD diet. While SAM levels in mice on the MD diet were comparable to those of mice on the control diet at ZT5 and ZT9, its levels at ZT13, ZT17, ZT21, and ZT1 were reduced by ∼50%. Furthermore, serum levels of MTA and SAH also exhibited circadian oscillation in mice fed the MD diet, but not in mice on the control diet (Figure 1E). Taken together, our findings suggested that dietary methionine regulated metabolic health by modulating circadian metabolism.

### Remodeling of hepatic rhythmic transcripts by MD diet

To investigate how the MD diet reshapes circadian metabolism, we conducted a circadian transcriptome analysis using liver tissues from mice after three weeks of being on either a control diet or an MD diet. Among the 4,375 oscillatory transcripts observed under the control diet, 2,841 shared rhythms with MD diet and 1,434 were cyclic only in the control diet. An additional 3,309 transcripts exhibited rhythmic behavior only during methionine deprivation (Figure 2A). Although the rhythmicity of core clock genes was preserved under both conditions, we observed that the expression of several genes was altered by the MD diet at specific time points (Figure S2A). For example, while the overall oscillatory expression patterns of *Per1* were similar between mice on the control and MD diets, its expression levels were upregulated at ZT13, ZT17, and ZT21 in mice on the MD diet (Figure S2B).

**Figure 2.**
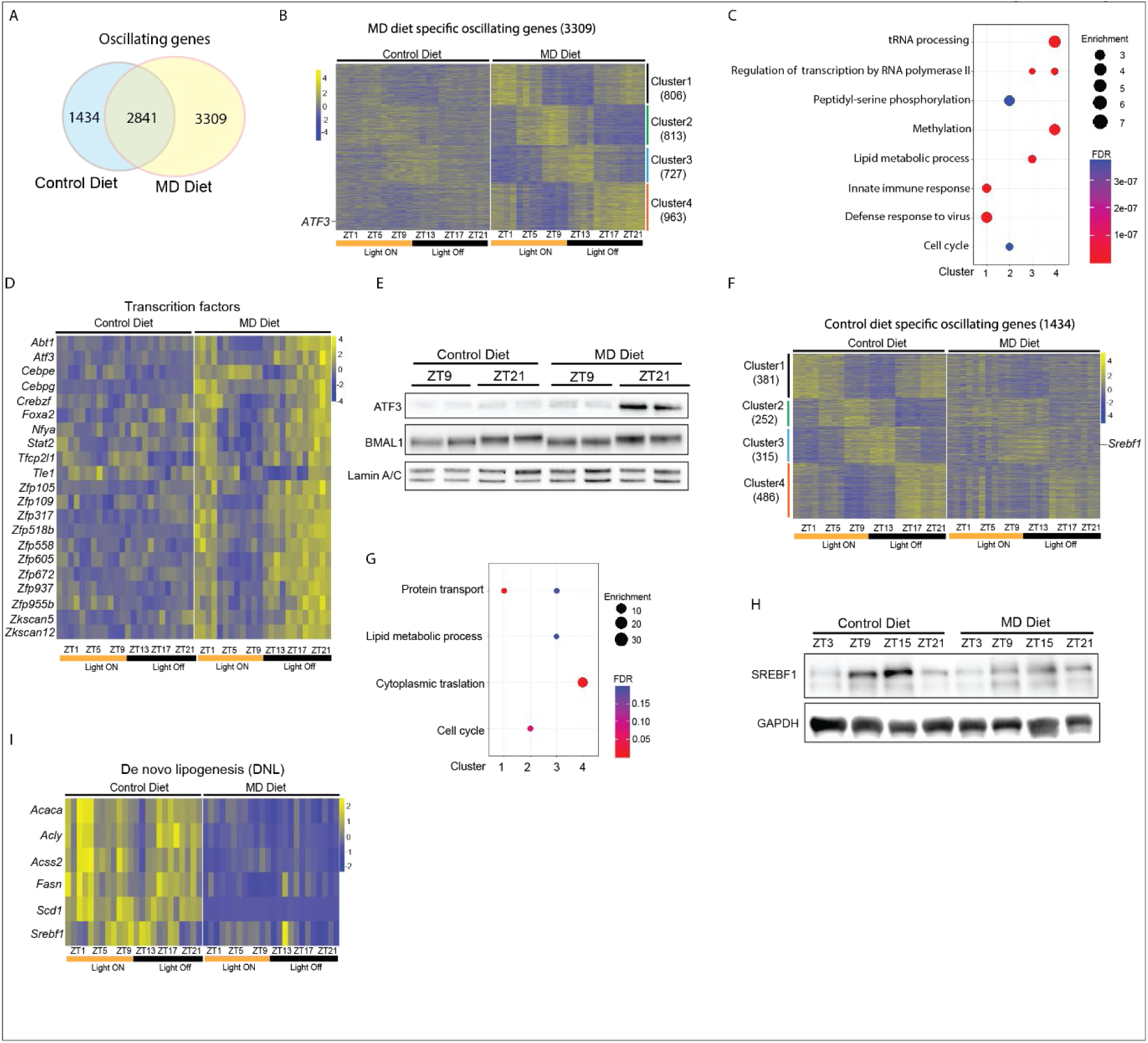
Reprogramming of Hepatic circadian transcriptome by Methionine deprivation. A. The Venn diagram illustrates the overlap of hepatic oscillating genes defined by MetaCycle in animals on control diet compared to those on the MD diet. B. Heatmap depicts the relative expression of 3309 genes which oscillate only in animals on MD diet. Genes are clustered in four groups based on clustering analysis by VSClust. Zeitgeber time for animal collection is indicated under the heat maps. Diet group is indicated above the heat maps, scale is indicated to the right of each heat map. At each time point, the heat maps include data from 4 animals per group. C. Gene ontology enrichment analysis of clusters 1, 2, 3, and 4 of MD diet specific oscillating genes. The color scale represents the FDR and Size of circle represents the enrichment. D. Heatmap depicts the relative expression of transcription factors with oscillating expression pattern only in animals on MD diet. E. The western blot depicts nuclear protein levels of ATF3 and BMAL1 at the indicated Zeitgeber time. Diet groups are indicated above the figure. Lamin A/C is included as loading controls. F. Heatmap depicts the relative expression of 1434 genes which oscillate only in animals on MD diet. Genes are clustered in four groups based on clustering analysis by VSClust. G. Gene ontology enrichment analysis of clusters 1, 2, 3, and 4 of control diet specific oscillating genes. The color scale represents the FDR and Size of circle represents the enrichment. H. The western blot depicts protein levels of SREBF1 at the indicated Zeitgeber time. Diet groups are indicated above the figure. GAPDH is included as loading controls. I. Heatmap depicts the relative expression of genes involved in de novo lipogenesis in animals. See also Figure S2.

By conducting fuzzy clustering analysis on genes exhibiting de novo oscillation in the liver under the MD diet, we found that these 3,309 genes were grouped into 4 distinct clusters, which were characterized as maximally expressed at ZT1 (cluster 1), ZT9 (cluster 2), ZT13 (cluster 3), and ZT21 (cluster 4) in mice fed on MD diet, respectively (Figure 2B). Gene ontology (GO) analysis revealed that 806 genes within cluster 1 were enriched in innate immune response and defense response to virus pathways (Figure 2C and S2C). Genes in cluster 2 (n=813) were related to cell cycle and peptidyl-serine phosphorylation pathways. Clusters 3 (n=727) showed enrichment of genes involved in lipid metabolism and regulation of transcription by RNA polymerase II pathways. Pathways were associated with 963 genes in cluster 4 including tRNA processing, methylation, and regulation of transcription by RNA polymerase II (Figure 2C and S2D). Notably, cluster 4 included several transcription factors, such as ATF3, FOXA2, and STAT2, which exhibited increased expression levels at ZT17, ZT21 and ZT1 in mice on MD diet but not in mice on control diet (Figure 2D). Consistent with the RNA-seq data, elevated ATF3 protein levels at ZT21 were observed only in mice on the MD diet, while BMAL1 levels remained comparable between the two diet groups (Figure 2E). Interestingly, both the transcription factor STAT2 and its target genes, which are involved in the anti-viral response, showed increased expression at ZT1 and ZT21 in mice fed the MD diet (Figure 2D & S2C). These findings suggest that the MD diet-induced de novo oscillation may be regulated by novel cyclic transcription factors in response to methionine starvation.

We found that 1,434 transcripts, which lost their oscillations of expression in mice placed on the MD diet, were grouped into four clusters (Figure 2F). Genes in Clusters 1 (n=381) and 2 (n=252) were enriched in protein transport and cell cycle pathway, respectively. Cluster 3 (n=315) showed the enrichment in protein transport and lipid metabolic process pathways. The 486 genes in cluster 4 were associated with cytoplasmic translation pathway (Figure 2G). We noted the oscillation of transcription factor SREBF1 in cluster 3 was lost in mice fed on MD diet (Figure 2F). Immunoblotting revealed that the rhythmicity of SREBF1 protein was also compromised by the MD diet (Figure 2H). Importantly, we found the expression levels of genes in de novo lipogenesis which are regulated by SREBF1 were down-regulated by MD diet (Figure 2I). On the other hand, we also noticed that the expression levels of *Ppara,* a key regulator in lipid metabolism, and genes involved in the lipid oxidation were decreased in mice on MD diet (Figure S2E &S2F). These findings imply that MD diet remodels the diurnal lipid metabolism via repressing both lipogenesis and lipid oxidation in liver.

### MD diet-induced de novo oscillations were entrained by the feeding cycle

Although feeding-fasting cycles have a significant impact on the oscillation of hepatic transcripts, behavioral rhythms driven by light-dark cycles also entrain hepatic circadian rhythms^30^. Time-restricted feeding (TRF) can decouple circadian oscillators in peripheral tissues from behavioral rhythms^31^. To further investigate how the MD diet influences hepatic rhythms, mice were placed on a time-restricted feeding (TRF) protocol for one week. In this protocol, food was available exclusively during either the dark or light phase (Figure 3A). Compared to mice fed on control diet, mice fed on MD diet displayed higher levels of circulating FGF21 only during the feeding phase (Figure S3A). This suggests that the oscillatory serum FGF21 induced by the MD diet is entrained by the feeding cycle, rather than the light-dark cycle.

**Figure 3.**
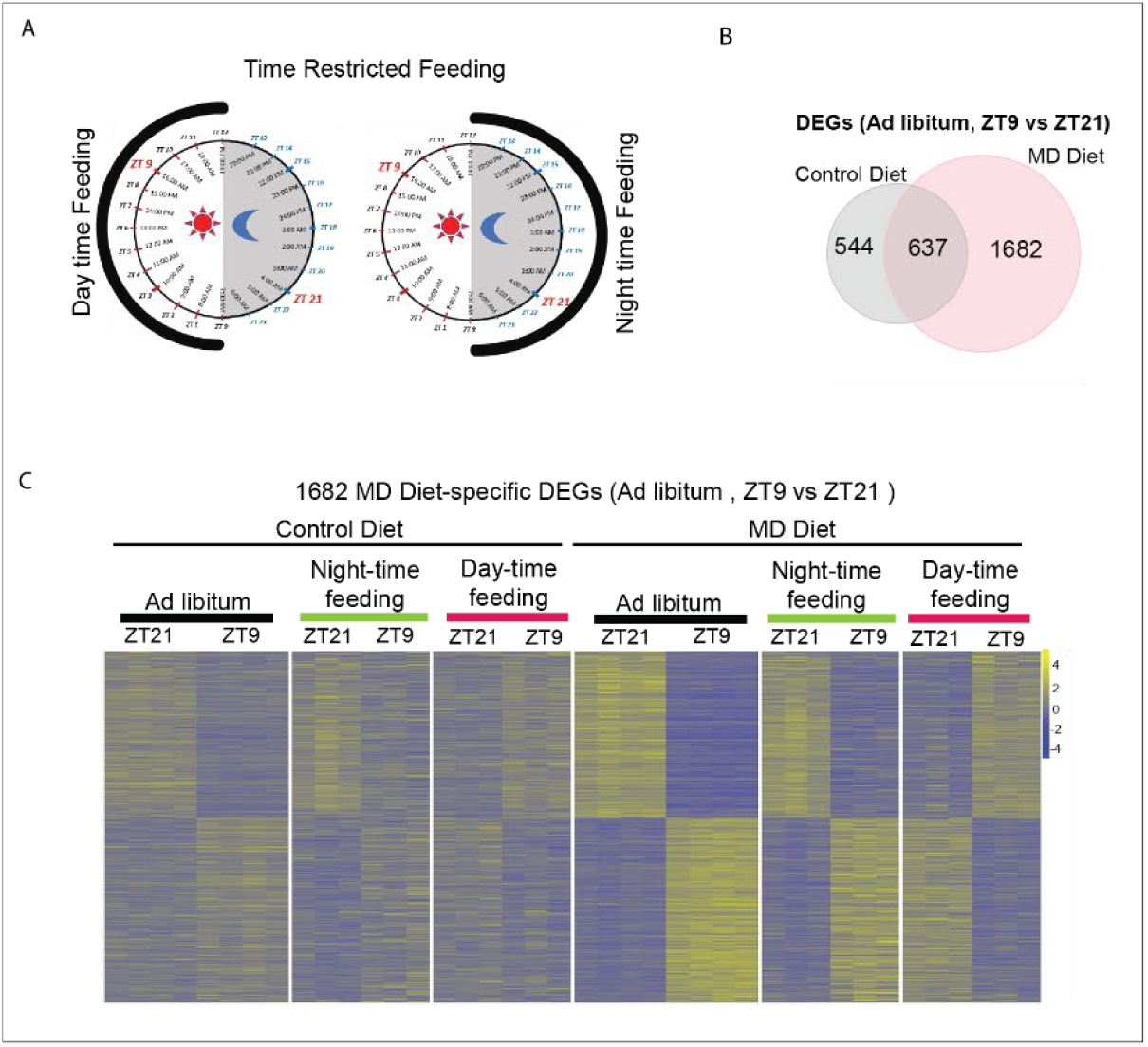
Reprogramming of hepatic transcripts entrained by feeding cycle. A. The schematic depicts the strategy employed for time restricted feeding. The mice were given access to Control diet or MD diet for a duration of 12 hour per day (either the dark phase or the light phase) for one week. B. Venn diagram showing overlap of differentially expressed genes (DEGs) (ZT9 vs ZT21, Ad libitum feeding for three weeks) between control diet and MD diet. The DEGs were determined by a combination of FDR < 0.05 and the absolute value of the log FC >0.7. C. Heatmaps depict relative expression of 1682 transcripts that are differentially expressed between ZT9 and ZT21 in MD diet only (ad libitum feeding) in mice subjected to time restricted feeding for one week and ad libitum feeding for three weeks. Feeding regimen and Zeitgeber time are indicated to the top of the heat map. See also Figure S3.

To investigate whether the MD diet-induced reprogramming of hepatic circadian rhythms is entrained by the feeding cycle, we first identified 1,682 genes that were differentially expressed between ZT9 and ZT21 in ad libitum-fed mice on the MD diet but not control diet, defining these as MD diet-specific differentially expressed genes (DEGs) (Figure 3B). We then examined the expression levels of these genes at ZT9 and ZT21 in mice subjected to TRF. The results revealed that nearly all 1,682 genes exhibited oscillatory expression patterns in mice fed during the night-time and day-time (Figure 3C). Importantly, the rhythmic expression of MD diet-specific DEGs was shifted in mice fed during the day-time compared to those fed ad libitum or during the night-time (Figure 3C). Specifically, cyclic genes involved in transcription regulation and tRNA processing, which were upregulated at ZT21 in both ad libitum- and night-time-fed MD diet mice, showed increased expression at ZT9 than ZT21 in mice fed during the day-time on the MD diet (Figure S3B & S3C). Taken together, these data demonstrate that the MD diet-induced de novo oscillation of transcripts is entrained by the feeding cycle.

### MD diet reshapes the circadian epigenetic landscape

Recent study demonstrates that five weeks on MD diet stimulates the loss of histone methylation in mouse liver^27^. To determine whether a shorter-term of MD diet feeding would influence histone modifications, we measured the levels of global histone methylations, including H3K4me3, H3K9me3, H3K36me3, and H3K27me3, in liver from mice on control and MD diet for three weeks.

Our data showed that these tri-methylation histone marks were intensely reduced by MD diet except for the repressive mark H3K27me3 (Figure 4A). Similarly, the methylation levels of H3K36me3, H3K4me3, and H3K9me3 also declined in HepG2 cells cultured with low methionine concentration, whereas H3K27me3 levels were not obviously reduced by methionine restriction *in vitro* (Figure 4B), suggesting that this mark is preserved in the face of limiting cellular methylation precursors.

**Figure 4.**
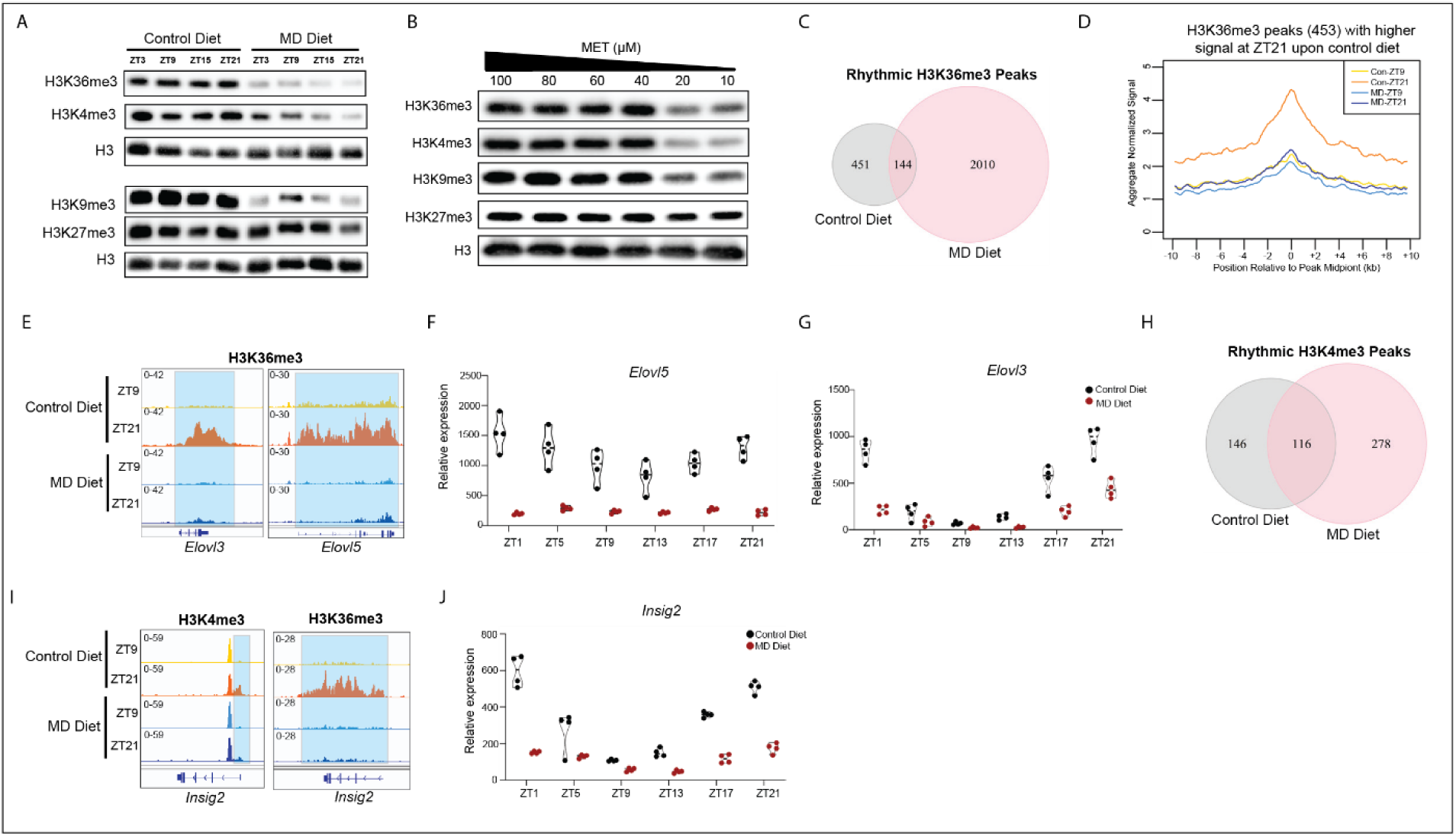
Methionine deprivation reshapes the rhythmic epigenetic landscape. A. Immunoblot of global H3 methylation levels in livers from mice fed on control and MD diet for three weeks. B. Immunoblot of H3 methylation levels in HepG2 cultured in medium with indicated methionine concentration for 24 hr. C. The Venn diagram depicts rhythmic H3K36me3 peaks in mice liver on control diet and MD diet measured by CUT&Tag. Rhythmic peaks were defined by comparing enrichment signals between ZT9 and ZT21 as described in Methods. D. Metagene profiles depicting H3K36me3 enrichment for 453 peaks with a stronger signal at ZT21 compared to ZT9 in control-diet fed mice (peak center ± 10 kb). E. IGV browser tracks showing H3K36me3 enrichment at *Elovl3* and *Elovl5* genes. Diet and Zeitgeber time for animal collection are indicated to the left, gene model is indicated below the data. F. TMM normalized expression values are presented in the graph for the *Elovl5* gene. G. TMM normalized expression values are presented in the graph for the *Elovl3* gene. H. The Venn diagram depicts rhythmic H3K4me3 peaks in mice liver on control diet and MD diet. Rhythmic peaks were defined by comparing enrichment signals between ZT9 and ZT21 as described in Methods. I. IGV browser tracks showing H3K4me3 and H3K36me3 enrichment at *Insig2* gene. Diet and Zeitgeber time for animal collection are indicated to the left, gene model is indicated below the data. J. TMM normalized expression values are presented in the graph for the *Insig2* gene. Diet group and Zeitgeber time are indicated on the graphs. See also Figure S4

Rhythmic histone methylation plays a crucial role in control of circadian clock^32,33^. We therefore questioned whether alterations in histone methylation are involved in MD diet-induced reprogramming of hepatic circadian rhythms. To detect the oscillation of histone modifications, we performed CUT&Tag for H3K4me3 and H3K36me3 marks using liver tissues, which were collected at ZT9 and ZT21, from mice upon control and MD diet. There are 595 oscillatory H3K36me3 peaks detected in control diet-fed mice. Strikingly, we found 2,154 rhythmic H3K36me3 peaks in mice upon MD diet (Figure S4A). Moreover, only 144 oscillatory H3K36me3 peaks were shared by mice in both conditions (Figure 4C). Metagene plot indicated that the rhythmic nature of H3K36me3 peaks specific to control diet was reflected in increased levels of modification at ZT21 compared with ZT9, whereas mice on MD diet failed to induce H3K36me3 abundance at ZT21 (Figure 4D). Examples of genes at which oscillations of H3K36me3 peaks over circadian time were undermined by MD diet included genes integral to the SREBF1 transcriptional program including the *Elovl3* and *Elovl5*, two key genes in lipid elongation (Figure 4E). Furthermore, the oscillation amplitude of these two genes was blunted in mice fed on MD diet (Figure 4F&4G). In contrast, the rhythmic nature of a larger peak set (1,373 H3K36me3 peaks) exhibited oscillation only in MD-fed mice, with failure of enrichment at ZT21 compared with ZT9. This group includes genes involved in bile acid and lipid metabolism such as *Nudt7* and *Ppara* (Figures S4B and S4C)^34^. We noted that the expression levels of *Nudt7* and *Ppara* were reduced by MD diet (Figure S2E&S4D). These data indicate that MD diet-altered rhythmic gene expression is associated with changes in cyclic H3K36me3 enrichment.

We next analyzed the oscillation of H3K4me3 and identified 262 and 394 oscillating H3K4me3 peaks in control and MD-diet fed mice, respectively (Figure S4E). Among of them, 116 peaks were rhythmic under both conditions (Figure 4H). There are 235 rhythmic H3K4me3 peaks with a higher enrichment and 159 peaks with a lower enrichment at ZT9 than ZT21 in MD diet-fed mice. Remarkably, the difference in the average signal of these peaks between ZT9 and ZT21 was more marked in MD diet-fed mice than mice upon control diet (Figure S4F), suggesting that methionine deprivation accentuates circadian alterations of this particular histone mark. We noted that the H3K4me3 peak at the lipogenic gene *Elovl2* promoter was specifically rhythmic in MD diet mice and this oscillatory H3K4me3 modification was associated with de novo oscillation of *Elovl2* expression upon MD diet (Figures S4G and S4H). Insulin-induced gene (INSIG)2 regulates SREBF1 function by inhibition of processing of SREBFs to their nuclear forms^35^. Two isoforms of *Insig2* mRNA transcript, *Insig-2a* and *Insig-2b*, are expressed in liver. While they encode identical proteins, their transcripts are regulated in a different way via alternative promoter usage^36^. Both the H3K4me3 and H3K36me3 modifications on *Insig2* were significantly rhythmic in mice on control diet whereas the oscillations of these histone marks were lost in MD diet-fed mice (Figure 4I). Correspondingly, we noted that the level and amplitude of *Insig2* mRNA oscillation was dramatically compromised by MD diet (Figure 4J). These data demonstrated that MD diet not only reduced the global histone methylation levels but also reshaped the rhythmic epigenetic landscape, leading to the reprogramming of circadian rhythms.

### MD diet altered the rhythmic GR binding to chromatin

The circulating FGF21 levels in mice were altered by the MD diet (Figure 1C). Previous studies have reported that FGF21 acts on the nervous system to increase systemic corticosterone levels^37,38^. Additionally, we found that the MD diet reduced *Insig2a* expression, which is induced by glucocorticoids via GR binding at the *Insig2* locus^39^. This led us to investigate whether the MD diet might affect the oscillation of circulating glucocorticoids and GR binding to chromatin.

We observed that serum glucocorticoid levels exhibited a rhythmic pattern in mice fed both the control and MD diets, peaking at ZT9, ZT13 and ZT17. Notably, the MD diet increased glucocorticoid levels at five of the six time points examined (Figure 5A). To determine whether the MD diet affects oscillatory GR binding to chromatin, we performed CUT&RUN analysis on liver samples collected at ZT9 and ZT21 from mice after three weeks on either the control or MD diet. GR enrichment was dramatically impacted by the MD diet, we identified 6,530 cyclic peaks in mice fed the control diet while there were only 3,707 cyclic GR peaks in mice fed the MD diet (Figure 5B). Of these, 1,715 oscillatory GR peaks were shared between the two conditions (Figure 5C), indicating that GR lost cyclic accumulation at 4,815 loci while gaining rhythmic behavior at approximately 2000. An exemplar gene with GR peaks exhibiting oscillation only under the control diet, *Insig2,* exhibited higher accumulation at ZT21 compared to ZT9. However, in mice fed the MD diet, GR binding to these loci did not show a significant change between ZT9 and ZT21 (Figure 5D). This loss of oscillatory GR binding in mice on the MD diet may explain the disrupted oscillation of *Insig2* mRNA observed in these mice (Figure 4H) and follows the general crippling regulation of lipid metabolism in these animals.

**Figure 5.**
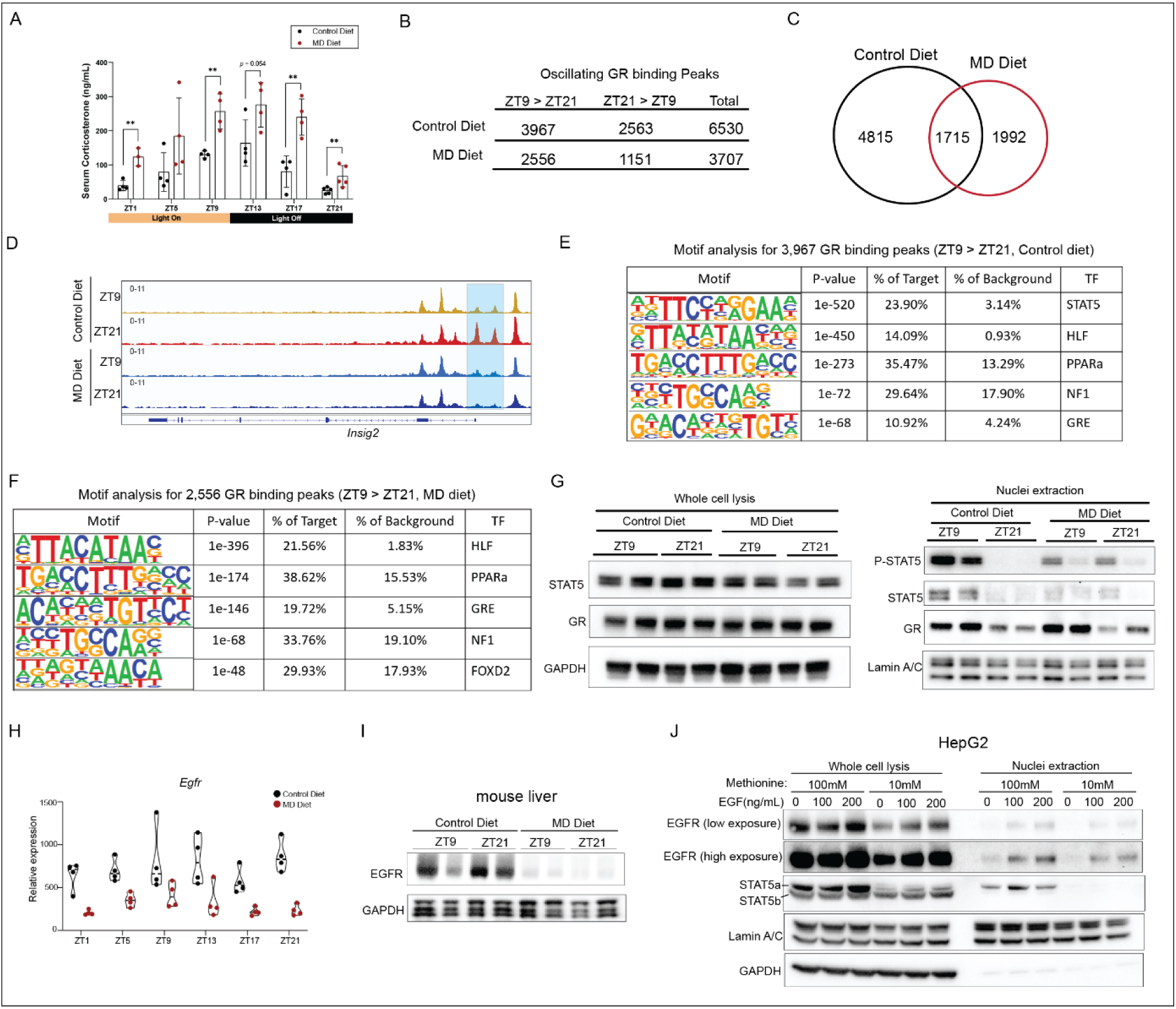
Methionine deprivation reshapes the circadian occupancy of glucocorticoid receptor (GR) A. Serum levels of corticosterone of six-month old male mice after three weeks on the indicated diet. The column graphs indicate mean value for 3-5 animals at indicated zeitgeber time points. Significance (**p<0.01, t test) is indicated with asterisks. B. The table shows the number of oscillating GR in liver from mice on indicated diet for three weeks. C. The Venn diagram depicts the rhythmic GR binding peaks in mice liver on control diet and MD diet detected by CUT&RUN. The rhythmic peaks were defined by comparing the enrichment signals between ZT9 and ZT21 as described in Methods. D. IGV browser tracks showing the GR enrichment at *Insig2* gene. Diet and Zeitgeber time for animal collection are indicated to the left. E. Motif analysis for 3,967 GR binding peaks with a stronger signal at ZT9 compared to ZT21 in control-diet fed mice. F. Motif analysis for 2,556 GR binding peaks with a stronger signal at ZT9 compared to ZT21 in MD diet fed mice. G. On the left, the western blot depicts the protein levels of STAT5 and GR in whole cell lysis of liver. GAPDH is included as the loading control. On the right, the western blot depicts the phosphorylation levels of STAT5, protein levels of STAT5 and GR in the nuclei extraction from liver. Lamin A/C is included as the loading control. H. TMM normalized expression values are presented in the graph for the *Egfr* gene. Diet group and Zeitgeber time are indicated on the graphs. I. The western blot depicts the protein levels of EGFR in liver from mice after three weeks on the indicated diet. GAPDH is included as the loading control. J. The western blot depicts the protein levels of EGFR and STAT5 in whole cell lysis and nuclei extraction of HepG2 cultured in medium with high methionine concentration (100 µM) or low methionine concentration (10 µM) for 40 hours following treated with EGF (0, 100 and 200 ng/mL) for 30 min. GAPDH and Lamin A/C are included as loading control. See also Figure S5

We found that GR exhibited higher occupancy at 3,967 peaks at ZT9 compared to ZT21 in control diet-fed mice (Figure 5B). De novo motif analyses for these peaks revealed that STAT5, HLF, PPARa, NF1 and glucocorticoid response elements (GRE) motifs were enriched (Figure 5E).

However, for 2,556 GR peaks that exhibited higher levels at ZT9 than ZT21 in mice fed on MD diet, the most significantly enriched motifs were HLF, PPARa, NF1 and GRE - the STAT5 motif was not detected. This suggests that the MD diet may alter oscillatory GR binding by modulating STAT5 signaling. Therefore, we sought to explore whether MD diet influenced the oscillation of STAT5 signaling. Immunoblotting results for whole-cell lysates indicated that STAT5 protein levels were modestly reduced in the liver of mice on the MD diet compared to those on the control diet at ZT21. In contrast, there was no difference in GR protein levels between the two conditions (Figure 5G). Surprisingly, we observed higher nuclear STAT5 levels and increased phosphorylation of STAT5 at ZT9 compared to ZT21 in the liver from mice on the control diet, whereas this oscillation was not observed in mice fed the MD diet. Consistent with higher circulating glucocorticoid levels at ZT9, nuclear GR levels were elevated at ZT9 compared to ZT21 in both control diet and MD diet groups (Figure 5G).

To further explore how the MD diet affects STAT5 signaling, we investigated the expression levels of primary receptors controlling STAT5 signaling and noted that the expression of *Egfr* (epidermal growth factor receptor) was reduced in mice on the MD diet (Figure 5H). Additionally, EGFR protein levels in the liver were dramatically reduced in these mice compared to those on the control diet (Figure 5I). To confirm the effects of methionine starvation on EGFR-STAT5 signaling, we exposed human hepatoma HepG2 cells to media with high (100 mM) or low (10 mM) methionine for 40 hours. We found that protein levels of EGFR and STAT5a were reduced upon exposure to low methionine. While EGF treatment increased EGFR internalization, nuclear STAT5 accumulation in the nucleus was only observed in cells grown in high concentrations of methionine (Figure 5J). These findings suggest that methionine starvation represses EGFR-STAT5 signaling and likely impacts STAT5/GR interactions on chromatin.

The protein-protein interaction between STAT5 and GR allows GR to bind to chromatin independently of the GRE^40^. The STAT5-GR interaction in hepatocytes plays a crucial role in regulating gene activation^41^. Catechol-O-methyltransferase (COMT) catalyzes the transfer of a methyl group from SAM to catecholamines^42^. We observed that GR showed higher occupancy at the promoter of one isoform of the *Comt* gene, which contains a STAT5 motif, particularly at ZT9 in mice on the control diet (Figure S5A). Interestingly, our RNA-seq data indicated that the expression of this isoform was repressed by the MD diet (Figure S5B). We also found that the MD diet reduced the expression of RNA transcripts and caused the loss of transcript oscillation (Figure S5C). Consistent with the reduced mRNA levels of *Comt*, we observed a decrease in COMT protein levels in mice on the MD diet. These results suggest that methionine deprivation may lead to a redistribution of SAM consumption through the regulation of STAT5-dependent cyclic GR binding, which controls COMT expression. These data show that the MD diet increases circulating glucocorticoid levels and reshapes the oscillatory GR binding, partly through the abolishment of STAT5 signaling oscillations.

### Altered oscillatory GR binding coupled with changes in H3K27ac enrichment

In response to GC stimulation, GR translocates into the nucleus and binds to tens of thousands of locations across the genome, predominantly at distal enhancers^43^. GR regulates gene expression by interacting with various epigenetic regulators^44^. To investigate whether oscillatory GR binding is associated with a cyclic epigenetic landscape, we performed CUT&Tag for H3K27ac, a marker for active enhancers and promoters, using liver samples collected at ZT9 and ZT21 from mice fed either a control or MD diet for three weeks. We found that GR peaks, which exhibited cyclic occupancy, showed similar oscillatory H3K27ac enrichment in mice on both diets (Figure 6A&6B). We noted that GR occupancy was positively associated with H3K27ac levels, suggesting that oscillatory GR binding may regulate the expression of target genes by modulating H3K27ac levels. For example, we observed that GR binding and H3K27ac were enhanced at the *Elovl5* loci at ZT21 compared to ZT9 in control diet-fed mice. However, no oscillatory binding was observed at this locus in MD diet-fed mice (Figure 6C). Consistent with the loss of oscillation in GR and H3K27ac binding, the oscillation of *Elovl5* transcript expression was also compromised in MD diet-fed mice (Figure 4F). We also observed cyclic GR binding and H3K27ac enrichment at the *Nudt7* gene loci in mice fed a control diet, whereas these oscillations were absent in mice fed an MD diet (Figure 6C). In contrast, de novo oscillations of GR binding and H3K27ac at the *Ppargc1a* and *Cyp39a1* genes were observed in mice on MD diet (Figure 4D). PGC-1α, encoded by the *Ppargc1a* gene, functions as a transcriptional coactivator and acts as a key link between the circadian clock and energy metabolism^45^. In mice fed on MD diet, both GR and H3K27ac enrichment were increased at *Ppargc1a* gene at ZT9 (Figure 6D). Meanwhile, the oscillation amplitude of *Ppargc1a* expression was elevated in MD diet-fed mice compared to control diet-fed mice (Figure 6E). *Cyp39a1* encodes oxysterol 7α-hydroxylase, an enzyme that converts 24-hydroxycholesterol into 7α-hydroxylated oxysterols, which are involved in bile acid metabolism^46^. Both GR and H3K27ac exhibited increased enrichment at the *Cyp39a1* loci at ZT9 specifically in mice fed the MD diet (Figure 6D). Notably, elevated *Cyp39a1* expression was detected in these mice (Figure 6E), suggesting that activation of *Cyp39a1* may be dependent on oscillatory GR binding. These findings demonstrate that the reshaped cyclic GR binding induced by the MD diet alters the oscillation of H3K27ac and gene expression.

**Figure 6.**
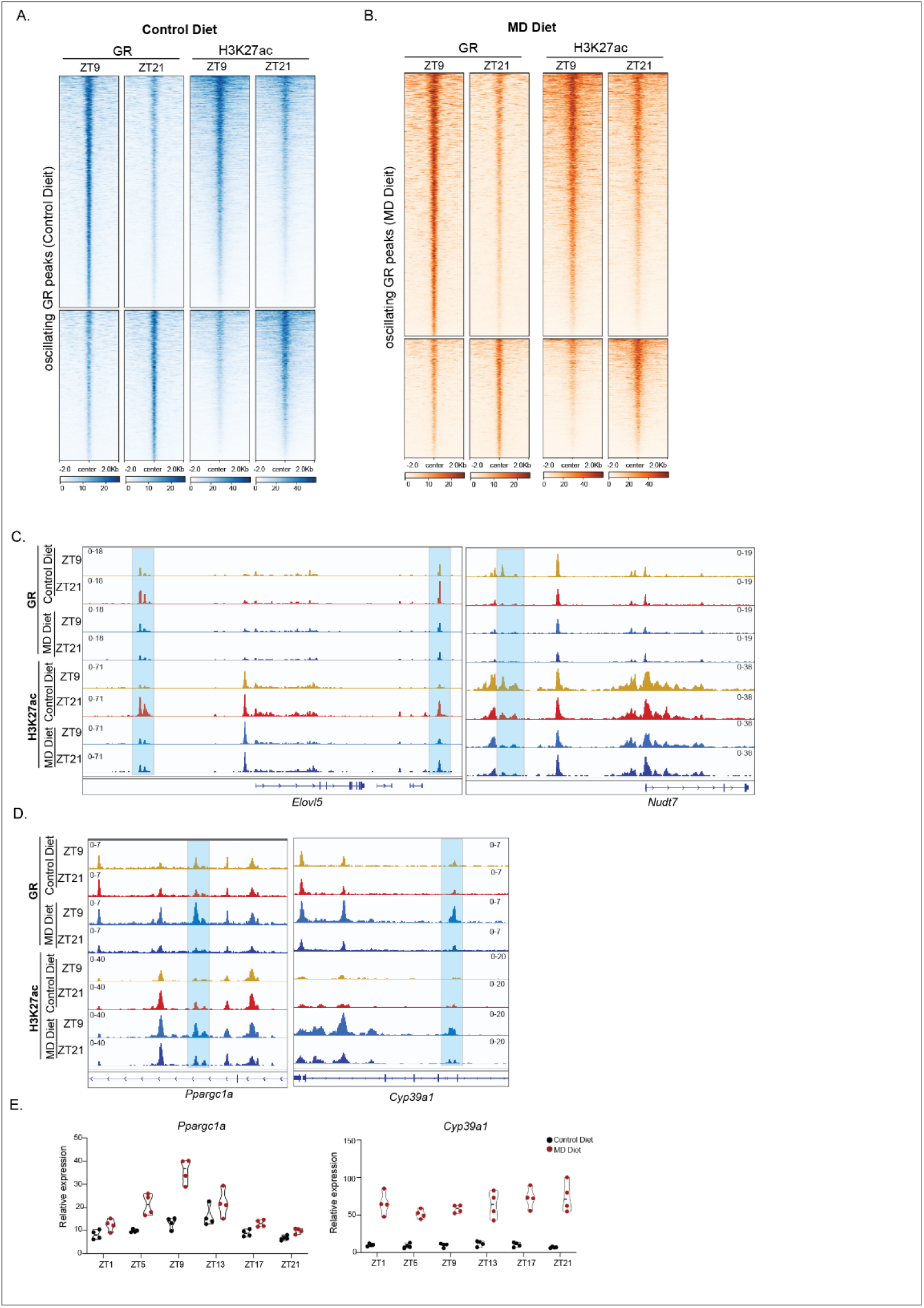
Oscillating GR binding associated with H3K27ac enrichment. A. The heatmaps depict the GR (left) and H3K27ac (right) enrichment levels at 6,530 rhythmic GR binding peaks at ZT9 and ZT21 in mice fed on control diet. B. The heatmaps depict the GR (left) and H3K27ac (right) enrichment levels at 3,707 rhythmic GR binding peaks at ZT9 and ZT21 in mice fed on MD diet. C. IGV browser tracks showing the GR and H3K27ac enrichment at *Elovl5* and *Nudt7* genes. Diet and Zeitgeber time for animal collection are indicated to the left. D. IGV browser tracks showing the GR and H3K27ac enrichment at *Ppargc1a* and *Cyp39a1* genes. Diet and Zeitgeber time for animal collection are indicated to the left. E. TMM normalized expression values are presented in the graph for the *Ppargc1a* and *Cyp39a1* genes. See also Figure S6 and S7

Next, we investigated how the abolished STAT5 signaling induced by the MD diet affected the cyclic epigenetic landscape. Specifically, we analyzed 948 GR binding peaks containing the STAT5 motif, which showed higher occupancy at ZT9 compared to ZT21 in mice on the control diet. Our results indicated that reduced GR binding to these STAT5 motifs was associated with decreased H3K27ac enrichment at ZT21 relative to ZT9. Notably, the difference in GR and H3K27ac enrichment between ZT9 and ZT21 was substantially diminished in mice on the MD diet (Figure S6A). Previous studies have shown that STAT5 regulates *Egfr* gene expression in the liver^47^. We found that both GR and H3K27ac were highly enriched at the *Egfr* locus at ZT9 in control diet-fed mice, correlating with higher *Egfr* expression in these mice (Figure 5H & S6B). Similarly, we observed that GR binding to STAT5 motifs at *Cyp7b1* and *Cyp2c70*, two genes involved in bile acid metabolism, was higher at ZT9 than at ZT21 in control diet-fed mice, accompanied by higher H3K27ac levels at ZT9 (Figure S6C). Additionally, we found altered oscillation patterns in the expression of *Cyp7b1* and *Cyp2c70* (Figure S6D), further suggesting that the MD diet may impact bile acid metabolism via modulating the GR binding.

Furthermore, we noted that the oscillation patterns of key genes involved in bile acid synthesis, including *Cyp7a1* and *Cyp27a1*, were altered by the MD diet (Figure S7A). Specifically, *Cyp7a1*, the rate-limiting enzyme in bile acid synthesis, showed increased expression at ZT17, ZT21, and ZT1 in MD diet-fed mice (Figure S7A). Consistent with the increased total bile acid levels in MD diet-fed mice (Figure S1F), we observed higher levels of primary, secondary, and conjugated bile acids in the serum of MD diet-fed mice compared to control diet-fed mice (Figure S7B). More importantly, we found that the levels of primary bile acids, including CA and α-MCA, and most conjugated bile acids in the liver were higher at ZT9 than at ZT21 in MD diet-fed mice, but not in control diet-fed mice (Figure S7C). These findings suggest that the MD diet reshapes the diurnal rhythms of bile acid metabolism.

These results demonstrate that the altered GR binding induced by the MD diet remodels the oscillations of gene expression and metabolism by modulating the cyclic epigenetic landscape.

### GR deletion blunts MD diet-Induced reprogramming of hepatic circadian rhythms

To validate whether GR mediates the reprogramming of circadian rhythms induced by the MD diet, we examined liver-specific GR knockout (LKO) mice on an MD diet. The LKO mice were generated by administering AAV8-TBG-Cre virus to *GR^fl/fl^* mice, with a control group receiving the virus without Cre recombinase. Two weeks post-virus injection, both control and LKO mice were placed on an MD diet for three weeks. Liver samples were collected at ZT9 and ZT21 (Figure 7A). Immunoblotting confirmed a significant reduction in GR protein levels in LKO livers compared to control livers (Figure 7B). Consistent with this decrease, RNA-seq data showed a reduction in the expression of the *Nr3c1* gene, which encodes the GR protein, at both ZT9 and ZT21 in LKO mice (Figures 7C & 7D). This further validates the successful deletion of GR in LKO livers. Notably, the *Cyp39a1* gene, which exhibits oscillatory binding by GR in response to the MD diet (Figure 6C), was among the top down-regulated genes following GR deletion at both ZT9 and ZT21 (Figures 7C & 7D). This suggests that GR is required for MD diet-induced activation of *Cyp39a1* gene expression in liver.

**Figure 7.**
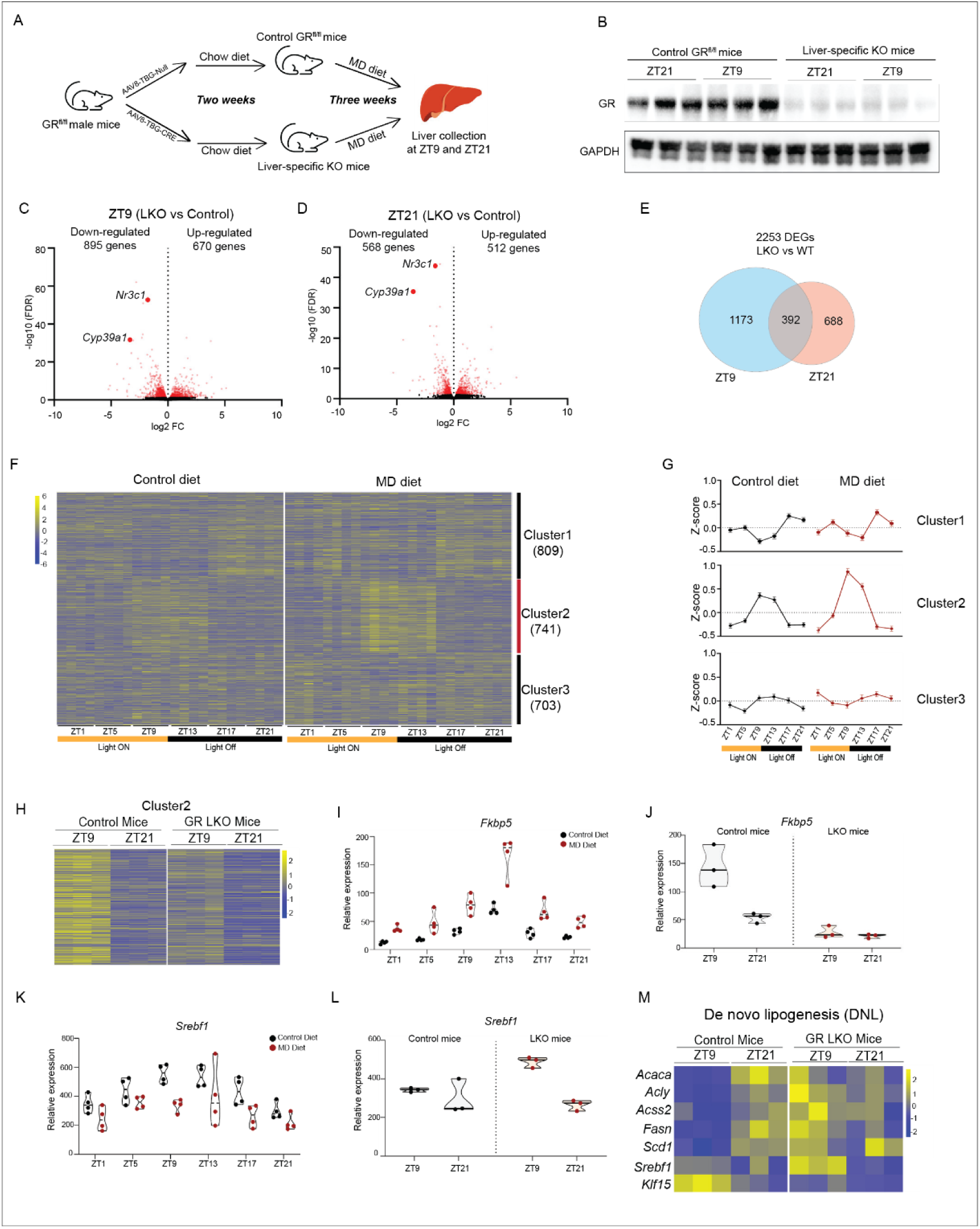
GR deletion in liver disrupts the diurnal rhythms of oscillating genes in mice fed on MD diet. A. The schematic depicts the experimental design for generating liver-specific GR KO (LKO) mice and control mice which were fed on MD diet for three weeks. B. The western blot depicts the protein levels of GR in whole cell lysis of liver from LKO and control mice on MD diet for three weeks. GAPDH is included as loading controls. C. Volcano plot showing the differential expressed genes (DEGs) in liver collected at ZT9 between LKO and control mice fed on MD diet for three weeks. DEGs are defined by FDR<0.05 and marked in red. D. Volcano plot showing the differential expressed genes (DEGs) in liver collected at ZT21 between LKO and control mice fed on MD diet for three weeks. DEGs are defined by FDR<0.05 and marked in red. E. The Venn diagram depicts DEGs in mice liver collected at ZT9 and ZT21 from LKO and control mice fed on MD diet. F. The heatmap displays combined DEGs showing significantly different expression between LKO and control mice at either ZT9 or ZT21. Genes are clustered into three groups based on clustering analysis using VSClust. G. The means and 95% confidence intervals of Z-score for Cluster 1, 2 and 3 of combined DEGs in liver from wild type mice on indicated diet for three weeks. H. The heatmap depicts the relative expression of cluster 2 genes at ZT9 and ZT21 in LKO and control mice fed on MD diet for three weeks. I. TMM normalized expression values of *Fkbp5* gene in liver from wild type mice fed on indicated diet for three weeks. J. TMM normalized expression values of *Fkbp5* gene in liver from GR LKO and control mice on MD diet for three weeks. K. TMM normalized expression values of *Srebf1* gene in liver from wild type mice fed on indicated diet for three weeks. L. TMM normalized expression values of *Srebf1* gene in liver from GR LKO and control mice on MD diet for three weeks. M. Heatmap depicts the relative expression of genes involved in de novo lipogenesis in GR LKO and control mice on MD diet for three weeks. See also Figure S8 and S9

RNA-seq analysis identified 1,565 differentially expressed genes (DEGs) in the livers of LKO versus control mice at ZT9 (Figure 7C). Among these, 895 down-regulated genes in LKO mice were enriched in lipid metabolism, cytoplasmic translation, and transcription regulation by RNA polymerase II, while 670 up-regulated genes were associated with immune system processes, lipid metabolism, and antigen processing (Figure 7C&S8A). At ZT21, a comparison between LKO and control mice identified 1,080 DEGs, with 568 down-regulated genes linked to lipid metabolism and cholesterol homeostasis, and 512 up-regulated genes enriched in immune system processes and lipid metabolism (Figure 7D&S8B). When comparing DEGs between ZT9 and ZT21, we found that 1,173 DEGs were specifically detected at ZT21, and 688 at ZT9. Only 392 DEGs were common at both time points, which implies that GR regulates distinct gene sets in a circadian-dependent manner on the MD diet.

We next examined the effect of GR deletion on genes exhibiting de novo oscillations in response to the MD diet. The expression of genes involved in transcription regulation and tRNA processing was comparable between LKO and control mice, and GR deletion did not affect the cyclic expression of most of these genes (Figure S9A&S9B). However, we observed that GR deletion led to a reduction in *Per1* expression at ZT9 compared to that of control mice, while other core clock genes remained unchanged (Figure S9C&S9D), suggesting that GR mediates the altered oscillation of *Per1* expression in response to the MD diet.

To further explore the role of GR in MD diet-induced reprogramming of hepatic circadian rhythms, we combined the DEGs identified at ZT9 and ZT21 between LKO and control mice and performed fuzzy clustering analysis on these 2,253 DEGs using RNA-seq data for detecting the oscillating genes. The genes were grouped into three clusters based on their oscillatory patterns (Figure 7F&7G). We noticed that genes in Cluster 2 exhibited an increased oscillation amplitude, with higher expression at ZT9 and ZT13 in MD diet-fed mice compared to control diet-fed mice (Figure 7F&7G). Interestingly, GR deletion in the liver reduced the expression levels of cluster 2 genes at ZT9 (Figure 7H). Among these genes, *Fkbp5*, a known GR target, showed a higher oscillation amplitude in MD diet-fed mice than in control diet-fed mice (Figure 7I). In control mice on the MD diet, *Fkbp5* expression was higher at ZT9 than at ZT21. In contrast, there was no difference in *Fkbp5* expression between ZT9 and ZT21 in LKO mice (Figure 7J). In response to MD diet, the gluconeogenic enzyme *Pck1*, another GR target gene, also showed an increased oscillation amplitude (Figure S9E). Similar to *Fkbp5*, the diurnal rhythm of *Pck1* expression was abolished by GR deletion in liver (Figure S9F). These findings demonstrate that GR mediates the increased oscillation amplitude of Cluster 2 genes induced by MD diet.

Fibroblast Growth Factor Receptor 1 (*Fgfr1*) serves as the receptor for FGF21^48^, and we observed that *Fgfr1* expression exhibited de novo oscillations in response to the MD diet (Figure S9G). While *Fgfr1* expression was higher at ZT9 than at ZT21 in control mice, this oscillation was lost in LKO mice (Figure S9H), suggesting that GR may mediate the MD diet-induced de novo oscillation of *Fgfr1*. KLF15, a fasting-induced transcription factor that represses *Srebf1* expression in the liver, has recently been shown to be regulated by GR^49,50^. We observed increased expression of *Klf15* at ZT21 in MD-fed mice compared to control-fed mice (Figure S9I). GR deletion resulted in the loss of oscillation in *Klf15* expression, with reduced *Klf15* levels observed at ZT9 in LKO mice compared to control mice (Figure S9J). Correspondingly, *Srebf1*, which lost its oscillation in mice fed on MD diet (Figure 7K), exhibited increased expression at ZT9 and showed a diurnal expression rhythm in LKO mice (Figure 7L). Moreover, we observed that genes involved in de novo lipogenesis were upregulated at ZT9 in LKO mice compared to control mice (Figure 7M), which further supports that GR- KLF15 pathway controls the loss of oscillation of *Srebf1* and repression of de novo lipogenesis in response to the MD diet. Taken together, these findings demonstrate that alterations in GR-dependent transcriptional regulation play a substantial role in MD diet-induced reprogramming of hepatic circadian rhythms.

## DISCUSSION

The circadian clock regulates various aspects of mammalian physiology, including metabolism and immune function^51^. It is well-established that disruptions to the circadian clock through nutritional challenges are closely linked to metabolic health^52^. While metabolic intermediates and enzymes involved in methionine metabolism have been shown to influence circadian gene transcription^53^, the role of dietary methionine in circadian homeostasis remains poorly understood. In the present study, we explore the effects of short-term dietary methionine deprivation on circadian rhythms in mice. Our findings demonstrate that dietary methionine governs the diurnal rhythms of circulating hormones, metabolites, the epigenetic landscape, transcription factor occupancy, and gene expression.

Dietary methionine restriction increases circulating FGF21, which regulates energy metabolism and remodels adipose tissue^20,21^. Consistent with this, our study detected elevated energy expenditure during the night in mice subjected to short-term methionine deprivation, likely linked to FGF21 oscillations induced by the methionine-deficient diet. We also found that the oscillations of circulating FGF21 driven by the methionine-deficient diet are synchronized with the feeding cycle, suggesting that FGF21 may be a key mediator in the de novo oscillations of hepatic gene expression triggered by methionine deprivation. However, there is ongoing debate about whether FGF21 signals directly to hepatocytes^54,55^. Some studies suggest that FGF21 regulates hepatic metabolism primarily through central actions in the brain^37,38^. Additionally, a recent study indicated that FGF21 does not mediate the effects of methionine restriction on hepatic lipid metabolism^21^. Although we observed MD diet-induced oscillations in the expression of its main receptor, FGFR1, its expression in the liver remains relatively low^56^. Therefore, the role of FGF21 in reprogramming hepatic circadian rhythms in response to methionine deprivation warrants further investigation in future studies.

While dietary methionine restriction increases circulating FGF21^21^, which activates the hypothalamic-pituitary-adrenal (HPA) axis and raises systemic glucocorticoid levels^37,38^, it remains unclear whether dietary methionine restriction directly affects glucocorticoids and their actions. Our study reveals that glucocorticoid levels are elevated under methionine deprivation (MD) diet. More importantly, we found that the MD diet significantly alters the rhythmic binding of its receptor, GR, in the liver. GR is known to play essential roles in both the circadian clock and metabolism^57–59^. Functional GREs have been identified in the regulatory region of the clock gene *Per1*, and GR binding activates *Per1* gene expression in response to glucocorticoids^59^. Our data show that the MD diet increases *Per1* expression at specific time points, and its rhythmic expression is disrupted in GR liver knockout (LKO) mice. These findings suggest that glucocorticoids and GR are key links between circadian rhythms and methionine metabolism. Recent studies have demonstrated rhythmic GR binding to chromatin in mice^58,60^, and Quagliarini et al. showed that nutritional challenges, such as a high-fat diet, can alter the GR binding pattern in the liver, enhancing GR-STAT5 co-occupancy at night^58^. In contrast, our data show that the MD diet represses the EGFR-STAT5 signaling pathway, abolishing STAT5 signaling oscillations. Furthermore, the rhythmic GR binding to STAT5 motifs is blunted by the MD diet, which in turn disrupts the oscillatory expression of target genes. These findings emphasize the role of the interaction between GR and STAT5 in the regulation of hepatic circadian rhythms in response to nutritional challenges. We also observed that EGFR expression is repressed during methionine starvation, which may mediate the disruption of STAT5 signaling due to methionine limitation. A recent study showed that methionine starvation leads to a decrease in the histone modification mark H3K79me2, resulting in lower STAT5 expression in T cells^61^. Our study further demonstrates the inhibitory role of methionine starvation on STAT5 signaling. Interestingly, it has also been reported that overexpression of FGF21 can blunt STAT5 signaling^62^. Thus, the molecular mechanisms underlying the inhibition of STAT5 signaling in response to methionine deprivation are likely complex.

Transcriptional rhythms are largely driven by the core circadian transcription machinery, including BMAL1, CLOCK, PERs, and CRYs, which are entrained by day/night cycles^54^. While we observed significant effects of the MD diet on cyclic gene expression, the oscillations of core clock genes (except for *Per1*) in the liver remained intact under methionine deprivation. These robust oscillations of the core clock machinery are also unaffected by other nutritional challenges, such as high-fat and ketogenic diets^8,10^. The high resilience of the core circadian clock to dietary composition may reflect its crucial role in maintaining metabolic homeostasis in response to feeding/fasting cycles^55^. Our study also reveals that several transcription factors (TFs), including ATF3, STAT2, and C/EBPγ, exhibit de novo oscillations upon methionine deprivation. These novel oscillating TFs may play a role in modulating circadian metabolism. Our data suggest that the MD diet recruits these non-core clock TFs as circadian regulators by modulating their transcription, leading to reprogramming of hepatic circadian rhythms in mice, including rhythms associated with lipid metabolism and immune response.

Accumulating studies have shown that methionine restriction affects histone methylation^22,25,27,61,63,64^. Our study demonstrates a trend where global histone methylation levels, including H3K36me3, H3K9me3, and H3K4me3, are positively correlated with serum SAM levels in MD-fed mice. Under MD diet conditions, histone methylation levels were further reduced during the dark phase, coinciding with lower SAM levels, supporting the idea that histones function as methyl-group sinks^13^. The steady-state levels of histone methylation are determined by the activity of histone methyltransferases, most of which have Km values for SAM in the 100 µM range, balanced by the activity of histone demethylases, some of which are known to be activated by methionine starvation through post-translational modifications^13,24^. Consistent with previous reports^64^, we did not observe large changes in H3K27me3 levels with methionine restriction.

Interestingly, the demethylases for H3K27me3 have distinct protein structures compared to those for other trimethylated histone residues, which may make their enzymatic activity less sensitive to SAM availability^65^. Further work will be needed to clarify the effects of methionine starvation on the modifications and enzymatic activity of H3K27me3 demethylases.

In addition to histones, methylation restriction has been shown to induce the demethylation of non-histone proteins^13,65^. For instance, methionine restriction activates the STING-TBK1-IRF3-type I interferon (IFN) pathway through the demethylation of cGAS^66^. Notably, we observed de novo oscillation in genes associated with the IFN I pathway, with higher expression in mice on the MD diet, suggesting that dietary methionine may reshape methylation oscillations in non-histone proteins. COMT, a methyltransferase that transfers methyl groups from SAM to catecholamines^42^, is repressed by the MD diet, providing insights into the molecular mechanisms underlying the redistribution of SAM consumption during methionine deprivation. Our findings highlight the impact of dietary methionine on various pathways, particularly through the modulation of methylation and demethylation processes.

Chronic methionine restriction remodels lipid and bile acid metabolism^18,26^. Here, we show that methionine starvation induces de novo oscillations in serum TG levels and bile acids in the liver, supporting the idea that methionine deprivation reshapes the diurnal rhythms of lipid and bile acid metabolism. Our data reveal that the MD diet regulates hepatic lipid metabolism by resetting oscillations of lipogenic genes at multiple levels. The master regulatory transcription factor SREBF1 loses rhythmic expression at the RNA level and is dramatically reduced at the protein level in response to the MD diet, potentially mediated by the GR-KLF15 pathway. Additionally, we demonstrate that the reshaped rhythmic transcription of lipid metabolism-related genes, such as *Insig2*, *Elovl3*, *Elovl5*, and *Elovl2*, is accompanied by altered rhythmic GR binding and histone modifications. Similarly, the MD diet also reshapes the rhythmic histone marks and GR occupancy at genes involved in bile acid metabolism, including *Nudt7*, *Cyp39a1*, *Cyp7b1*, and *Cyp2c70*. Notably, we show that oscillatory GR binding is required for the activation of *Cyp39a1* expression upon the MD diet. These findings suggest that an MD diet regulates the circadian rhythms of lipid and bile acid metabolism through alterations in circadian epigenetic dynamics and transcription factor binding, further highlighting the intricate connection between cellular metabolism, epigenetic regulation, and circadian rhythms.

### Limitations of Study

In our study, we highlighted the role of dietary methionine in control of circadian rhythms via modulation of methylation cycle. However, methionine availability has been shown to regulate various signaling pathways, including mTORC1 and AMPK (AMP-activated protein kinase) pathways^18,67^, which modulate the circadian clock in response to nutrient signals^68^. The precise functions of these pathways in the reprogramming of circadian rhythms by MD diet require further investigations.

## ACKNOWLEDGMENTS

We gratefully acknowledge the Epigenomics and DNA Sequencing Core Facility at NIEHS for their assistance with NGS library sequencing. We thank Greg Travlos and Ralph Wilson of the Clinical Pathology Group at NIEHS for their help with serum chemistry analysis. We also thank David Goulding and Terry Blankenship-Paris of the Comparative Medicine Branch at NIEHS for their support with the animal experiments. This work was supported by the Intramural Research Program of the National Institute of Environmental Health Sciences, NIH (ES101965 to P.A.W., ES090057 to J.A.C., ES103005 to L.J.D.).

## AUTHOR CONTRIBUTIONS

Y.L. and P.A.W. conceived the project and designed experiments. Y.L. performed experiments. Y.L. and S.A.G analyzed data. F.B.L. and L.J.D measured metabolites and bile acids. J.A.C. and R.H.O. provided the *GR^fl/fl^* mice. Y.L. and P.A.W. co-wrote the manuscript with input from all authors.

## STAR METHODS

## RESOURCE AVAILABILITY

### Lead contact

Further information and requests for resources and reagents should be directed to and will be fulfilled by the lead contact, Paul A. Wade.

### Materials availability

This study did not generate new unique reagents.

### Data and code availability

- All sequencing data have been deposited the BioProject of NCBI with accession number : GSE240881
- This paper does not report original code.
- Any additional information required to reanalyze the data reported in this paper is available from the lead contact upon request.

## EXPERIMENTAL MODEL AND SUBJECT DETAILS

### Animals

Six-month-old male C57BL/6J mice (JAX, 00064) were purchased from the Jackson Laboratory. Mice were raised under a 12-hour light:12-hour dark schedule in the specific-pathogen-free (SPF) conditions for 2 weeks to adapt to the housing condition. Mice were fed ad libitum either amino acid defined control diet (Envigo, TD.01084, 0.82% methionine) or methionine deficient diet (MD, Envigo, TD.140119, 0% methionine) for 1-3 weeks. For time-restricted feeding study, mice were allowed access to the food for 12 hour per day either during the dark phase or the light phase for one week. ZT0 (6 am, lights on) represents the beginning of the light phase, and ZT12 (6 pm, lights off) corresponds to the end of the light phase. Two-month-old *GR^fl/fl^* male mice, previously described^69,70^, were administered AAV8-TBG-Cre or AAV8-TBG-Control virus via retro-orbital injection to generate the GR liver-specific knockout (LKO) mice and control mice. All animal procedures were approved by the NIEHS Animal Care and Use Committee, and they were performed according to NIH guidelines for care and use of laboratory animals.

### Cell line

Human liver cancer cell HepG2 was obtained from ATCC and maintained in RPMI-1640 (GIBCO, 11875119) supplemented with 10% FBS at 37 °C with 5% CO_2_. For methionine starvation experiment, plated cells were washed with ice cold phosphate-buffered saline (Calcium & Magnesium free, PBS-CMF) twice and then cultured in the RPMI Medium (No Methionine, GIBCO, A1451701) supplemented with 10% dialyzed fetal calf serum (Thermo Fisher Scientific, A3382001) and varying L-Methionine concentration.

## METHOD DETAILS

### Serum chemistry tests

Blood was collected from mouse heart and placed at room temperature for 1 hour for clotting. After a centrifugation at 3,000 rpm for 10 min at 4 °C, the serum was transferred to new tube and stored at -80°C. Serum lipid panel (Cholesterol, HDL Cholesterol, LDL Cholesterol and Triglycerides) and liver panel (AST, ALT, ALP, SDH and Total Bile Acids) were measured by Clinical Pathology Laboratory in NIEHS.

### Indirect calorimetry

The energy metabolism was assessed using the TSE phenoMaster System (TSE Systems) in NIEHS. Mice were fed on control diet and methionine deficient diet for 7 days and then transferred into CaloCages for 3 days with unlimited access to food and water. CO_2_ production and O_2_ consumption were measured, and energy expenditure was calculated using the following equation: total energy expenditure (kcal/h) = 3.941 × VO_2_ (l/h) + 1.106 × VCO_2_ (l/h). The first 24h were defined as acclimation period and excluded from data analysis. Data were normalized by body weight.

### Glucose and insulin tolerance tests

For glucose tolerance test, mice were fasted overnight for 14 hours after fed on control diet and Methionine deficient diet for 2 weeks. Glucose (2 g/kg) was injected intraperitoneally and then blood glucose was measured using a glucometer at 0, 30, 60, 90, and 120 min. Intraperitoneal insulin tolerance tests (IPITTs) were performed by withholding food from mice for 4 hours. Insulin (0.8 U/kg) was injected intraperitoneally and blood glucose measurement was taken with a glucometer at 0, 30, 60, 90, and 120 min.

### Serum Metabolites measurement

Serum proteins were precipitated with 3 volumes of 90% acetonitrile containing 10 mM ammonium acetate pH 5.3. The supernatant was evaporated under vacuum and reconstituted in 10 mM ammonium acetate pH 5.3 for analysis. Methionine (MET), S-adenosyl-methionine (SAM), S-adenosyl-homocysteine (SAH) and methylthioadenosine (MTA) were separated by reverse phase liquid chromatography on a Raptor Fluorophenyl column (Restek). Analytes were detected using tandem quadrupole mass spectrometry. Relative abundance was calculated by averaged peak area of three instrumental injections.

### Serum FGF21 and glucocorticoids measurement

Blood was collected from the mouse heart and allowed to clot at room temperature for 1 hour. After centrifugation at 3,000 rpm for 10 minutes at 4°C, the serum was transferred to a new tube and stored at -80°C. The concentration of FGF21 in the serum was measured using the Mouse/Rat FGF-21 Quantikine ELISA Kit (R&D Systems, MF2100) according to the manufacturer’s instructions. Serum corticosterone levels were assessed using the Corticosterone Multi-Format ELISA Kit (Arbor Assays, K014-H1) following the provided protocol.

### Body composition

Body composition (fat and lean mass) was determined *in vivo* with the Bruker minispec LF90II

### Bile acid in liver and serum

Proteins were precipitated from 20 µL serum with 70 µL of acetonitrile and 10 µL isotope labeled internal standard solution in acetonitrile. After vortex mixing and centrifugation, the supernatant was collected for analysis. Liver samples of approximately 20 mg were homogenized with a bead mill in 480 µL acetonitrile per 20 mg tissue. 125 µL aliquots of the supernatant were removed and spiked with isotope labeled internal standard solution prior to drying by vacuum centrifugation. Liver samples were reconstituted in 120 µL 75% acetonitrile prior to analysis. Unconjugated bile acids and taurine conjugated bile acids were separated by reverse phase gradient liquid chromatography on an ACE Excel CN-ES column. The injection volume was 2 µL. Analytes were detected using negative ion electrospray tandem quadrupole mass spectrometry on a Sciex 7500 instrument. The [M-H]^-^ ion was isolated in Q1 for all analytes. Unconjugated bile acids were detected as the unfragmented ion after collision induced dissociation at -40 eV. Taurine conjugates were fragmented at -130 eV and the *m/z* -80.0 fragment monitored. Calibration curves were built in charcoal stripped serum.

### RNA-seq and data analysis

Total RNA was extracted from snap-frozen liver with TRIzol Reagent (Invitrogen, 15596026). After rRNA depletion with Ribo-Zero H/M/R Gold rRNA Removal Kit (Illumina, MRZG126), 250 ng of purified RNA was used for RNA-seq library preparation with TruSeq Stranded Total RNA Library Prep Kit. Sequencing was Illumina NextSeq 500 or Illumina NovaSeq 6000 (single-end, 75 bp). Reads were mapped to mm10 reference genome with STAR v2.7.9a at default parameters^71^, and then DEGs were identified by DESeq2 v1.32.0^72^.

Rhythmicity of gene expression were detected by R package Metacycle v1.2.0 using incorporated JTK methods^73,74^. Rhythmic genes were defined as transcripts with adjusted p value less than 0.005 with a window of 20-24 hours for the determination of circadian periodicity. Fuzzy Clustering analysis was conducted with R package VSClust v1.1.0^75^. Gene ontology enrichment tests were performed on the DAVID Bioinformatics Resource website (https://david.ncifcrf.gov/)^76^.

### CUT&Tag

The cleavage under targets and tagmentation (CUT&Tag) technology was performed as reported with minor modifications^77^. Briefly, Nuclei were extracted from snap-frozen tissue with single nucleus isolation kit (Invent Biotechnology, SN-047) following the manual of kit. After lightly fixed with 0.1% formaldehyde for 2 min at room temperature, about 50,000 nuclei were used. Antibodies against H3K4me3 (Active Motif, 39159), H3K27ac (Invitrogen, MA5-23516) and H3K36me3 (Abcam, ab9050) were used as primary antibodies at 1:100 dilution. Tagmented DNA was purified with Qiagen MinElute PCR purification kit (QIAGEN, 28004) and amplified with High-fidelity PCR mix (NEB, M0541S). Libraries were purified with AMPure XP beads (Beckman, A63881) and sequenced on Illumina NextSeq 500 or Illumina MiSeq instruments.

### CUT&Tag data analysis

Raw read pairs were adapter-trimmed by Cutadapt v1.2 with parameters "--quality 20 --nextera --stringency 5 --length 20 --paired"^78^, then mapped to the mm10 reference genome by Bowtie v2.1 with parameters "-X 2000 --fr --local --sensitive-local"^79^. Alignments were filtered to retain only proper pairs with MAPQ at least 5 via samtools (-q 5 -f 2). Duplicate fragments were removed with MarkDuplicates.jar (REMOVE_DUPLICATES=TRUE) from the Picard tool suite v1.110. For samples with multiple libraries, alignment files were concatenated with MergeSamFiles.jar from the Picard tool suite v1.110. Peak calls for each sample were made by HOMER v4.10.3 with parameters "-region -size 1000 -minDist 2500 -L 0" for H3K36me3 and "-region -size 500 -minDist 1000 -L 0" for H3K4me3, followed by filtering out calls with overlap to mm10 blacklist regions^80^. For each histone mark, a set of unified peaks was defined as regions called as peaks in at least two samples; unified peaks are limited to canonical chromosomes, neighboring regions within 50bp are merged, and a minimum peak size of 50bp is applied. Oscillating peaks were identified with EdgeR v3.30.3 at FDR 0.05^81^, based on fragment-per-peak counts collected by Bedtools v2.29.2 function ‘coverage’ with the ‘-counts’ option^82^.

### CUT&RUN

CUT&RUN was performed according to a previously reported protocol with minor modifications^83^. Briefly, frozen liver tissues were homogenized in PBS-CMF containing 0.4% formaldehyde, followed by a 5-minute incubation at room temperature. Nuclei were isolated using a nuclei extraction buffer (20 mM HEPES-KOH pH 7.9, 10 mM KCl, 20% glycerol, 0.1% Triton X-100, 0.5 mM spermidine, and EDTA-free protease inhibitor), and approximately 300,000 nuclei were used for the experiment. The bead-nuclei slurry was incubated with antibodies against GR (Proteintech, 24050-AP, 1:50 dilution) or with no primary antibody as a negative control. After two washes with wash buffer (20 mM HEPES pH 7.5, 150 mM NaCl, 0.5 mM spermidine, and EDTA-free protease inhibitor), the bead-nuclei slurry was incubated with a secondary antibody (Guinea pig polyclonal anti-Rabbit IgG, Antibodies Online, ABIN101961) at a 1:100 dilution. The slurry was then incubated with pA-MNase (gift from Henikoff Lab, 1:100 dilution) for 1 hour at 4 °C. After two additional washes with wash buffer, beads were resuspended in ice-cold low-salt rinse buffer (20 mM HEPES pH 7.5, 0.5 mM spermidine). After removing the liquid, ice-cold incubation buffer (3.5 mM HEPES pH 7.5, CaCl_2_) was added, and the slurry was incubated on ice for 10 minutes. Following liquid removal, stop buffer (170 mM NaCl, 20 mM EGTA) was added, and the beads were incubated at 37 °C for 25 minutes. DNA was extracted using phenol-chloroform. Libraries were prepared using the NEBNext® Ultra™ II DNA Library Prep Kit (NEB, E7645L) with 10 ng of DNA as input, purified using AMPure XP beads (Beckman, A63881), and sequenced on an Illumina NextSeq 500 instrument.

### CUT&RUN data analysis

Raw read pairs were aligned to the mm10 reference genome using Bowtie v2.1 with the parameters "-X 2000 --local --very-sensitive"^79^. Alignments were filtered to retain only properly paired reads with a MAPQ score of at least 5 using samtools (-q 5 -f 2). Duplicate fragments were removed with MarkDuplicates.jar (REMOVE_DUPLICATES=TRUE) from the Picard tool suite v1.110. Peak calling for each sample was performed with MACS3 v3.0.0b3 using the parameter "-q 0.0001"^84^. A set of unified peaks was defined as regions called as peaks in at least two samples. These unified peaks were limited to canonical chromosomes, with neighboring regions within 50 bp merged and a minimum peak size of 50 bp applied. Calls overlapping with mm10 blacklist regions were excluded. Oscillating peaks were identified using EdgeR v3.30.3 with a FDR threshold of 0.05^81^, based on fragment-per-peak counts collected by featureCounts v2.0.6 with the ‘--countReadPairs’ option^85^. Motif analysis was performed using HOMER v4.10.3^80^.

### Protein extraction and Immunoblotting

For total protein extraction, frozen tissues were pulverized on dry ice and cells were lysed in RIPA buffer (Thermo Fisher Scientific, 89901) containing Halt Protease and Phosphatase Inhibitor Cocktail (Thermo Fisher Scientific, 1861280). Lysates were placed on the ice for 30 min followed by incubation at 4 °C for 90 min on a rotator. Following a 15-min centrifugation at 14,000 g at 4 °C, supernatant was collected.

For collection of the nuclear extracts from liver tissues, pulverized frozen tissues were homogenized in 1 mL of homogenization buffer (250 mM Sucrose, 25 mM KCl, 10 mM MgCl2, 20 mM Tricine-KOH pH 7.8, 1 mM DTT, 0.5 mM spermidine, 0.15 mM spermine, 0.3% NP-40, Halt Protease and Phosphatase Inhibitor Cocktail) with Dounce tight pestle for 25 strokes and then incubated on ice for 10 min. After centrifugation at 500g for 5 min at 4 °C, the nucleus pellet was resuspended in RIPA buffer (Thermo Fisher Scientific, 89901), supplemented with protease inhibitor and phosphatase inhibitor (Thermo Fisher Scientific, 1861280), and incubated on ice for 1 hr. After centrifugation at 14,000 g for 10 min, the supernatant was recovered as soluble nuclear extracts. Histone proteins enriched lysates were prepared with histone extraction kit following the instruction manual (Active Motif, 40028). Proteins were separated on 4-12% Bis-Tris Nupage gels (Invitrogen, NP0336BOX) and then transferred onto PVDF membrane (Bio-Rad, 162-0218) followed by incubation with Primary antibodies against ATF3 (Cell Signaling, 33593), LAMIN A/C (Cell Signaling, 4777), H3K36me3 (Abcam, ab9050), H3 (Cell Signaling, 9715), H3K9me3 (Abcam, ab8898), H3K4me3 (Millipore Sigma, 04-745), H3K27me3 (Millipore Sigma, 07-449), GAPDH (Santa Cruz, sc-32233), BMAL1 (Cell Signaling, 14020), GR (Proteintech, 24050-AP), COMT (Invitrogen, PA5-95276), STAT5 (Cell Signaling, 94205) , STAT5A/B (Proteintech, 13179-AP), EGFR (Proteintech, 30847-AP), Phospho-STAT5 (Cell signaling, 9359) and SREBF1 (Abcam, ab3259). Blots were visualized using the Chemidoc XRS system (Bio-Rad).

## QUANTIFICATION AND STATISTICAL ANALYSIS

Statistical analysis was performed using GraphPad Prism 9 software. Detailed descriptions of sample numbers and statistical tests can be found in figure legends. Statistical significance was determined by unpaired t-test, or one-way analysis of variance (one-way ANOVA) and two-way analysis of variance (two-way ANOVA) followed by post hoc Tukey’s tests. p < 0.05 was considered as statistically significant.

