## Supplemental Figures 1-9 for "Methionine Deprivation-induced Reprogramming of Hepatic Rhythms Is Mediated by Glucocorticoid Receptor"

**Supplementary Information**

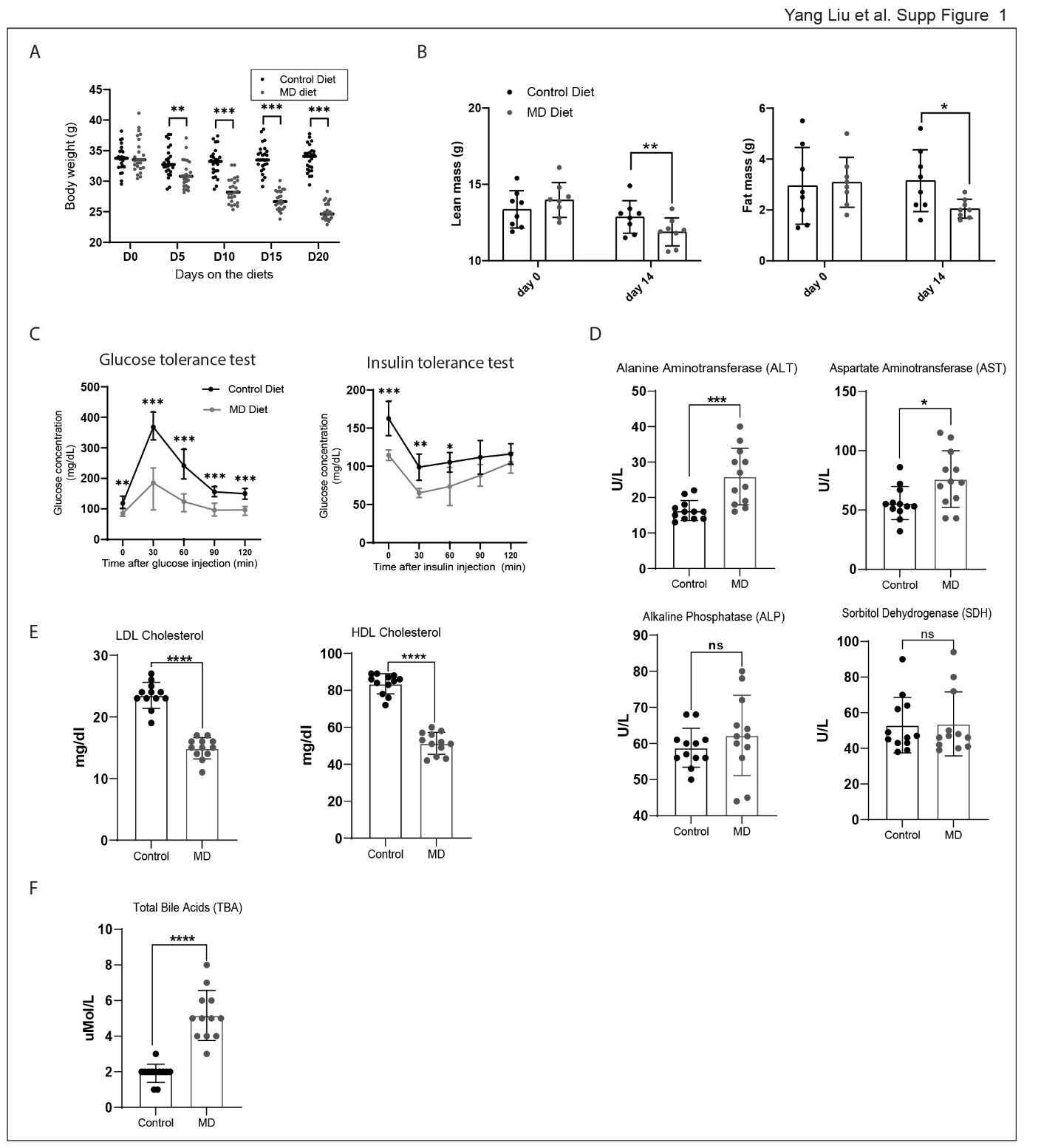

**Figure S1. Short-term Methionine deprivation affects metabolic health, related to Figure 1**

A. The scatter plot depicts body weights of individual mice fed as a function of time on diet. Mean value for each group is indicated by a horizontal black line. Significance between groups is indicated with asterisks (**p<0.01,***p<0.001, unpaired t tests).
B. Lean (left panel) and fat (right panel) mass in mice was measured before and after 2 weeks on the indicated diet as described in Methods. Significance between groups is indicated with asterisks (*p<0.05,**p<0.01, unpaired t tests).
C. Glucose tolerance tests (GTT) and insulin tolerance tests (ITT) were performed on mice after two weeks on the indicated diets. Significance between groups is indicated with asterisks (*p<0.05,**p<0.01,***p<0.001, unpaired t tests).
D. Serum levels of the indicated liver marker enzymes were determined as described in Methods. The column graph indicates mean (column height), standard deviation, as well as values from individual animals. Significance between groups is indicated with asterisks (*p<0.05,***p<0.001, unpaired t tests).
E. Serum levels of LDL and HDL cholesterol were determined as described in Methods. The column graph indicates mean (column height), standard deviation, as well as values from individual animals. Significance between groups is indicated with asterisks (****p<0.0001, unpaired t tests).
F. Serum levels total bile acids were determined as described in Methods. The column graph indicates mean (column height), standard deviation, as well as values from individual animals. Significance between groups is indicated with asterisks (****p<0.0001, unpaired t tests)

**Figure S2. Methionine deprivation reshapes the oscillation of genes and pathways in liver, related to Figure 2**

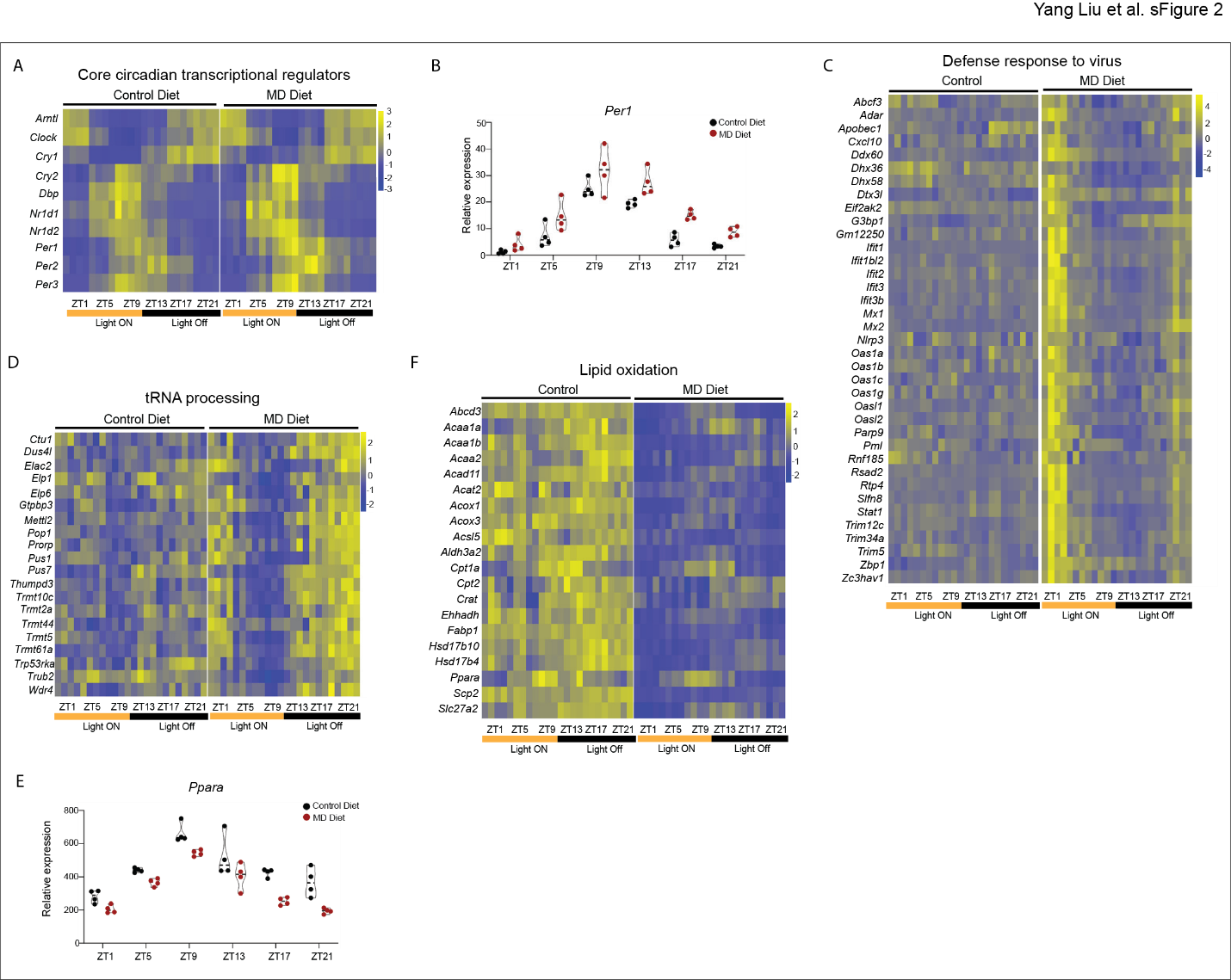

A. Heatmap depicts the relative expression of core circadian clock genes in animals on both control and MD diet. Zeitgeber time for animal collection is indicated under the heat maps. Diet group is indicated above the heat maps, scale is indicated to the right of each heat map. At each time point, the heat maps include data from 4 animals per group.
B. TMM normalized expression values are presented in the graph for the *Per1* gene. Diet group and Zeitgeber time are indicated on the graphs.
C. Heatmap depicts the relative expression of genes related to defense response to virus in animals.
D. Heatmap depicts the relative expression of genes enriched in tRNA processing in animals.
E. TMM normalized expression values are presented in the graph for the *Ppara* gene. Diet group and Zeitgeber time are indicated on the graphs.
F. Heatmap depicts the relative expression of genes involved in lipid oxidation in animals.

**Figure S3. MD-diet induced de novo oscillation of pathways entrained by time-restricted feeding, related to Figure 3**

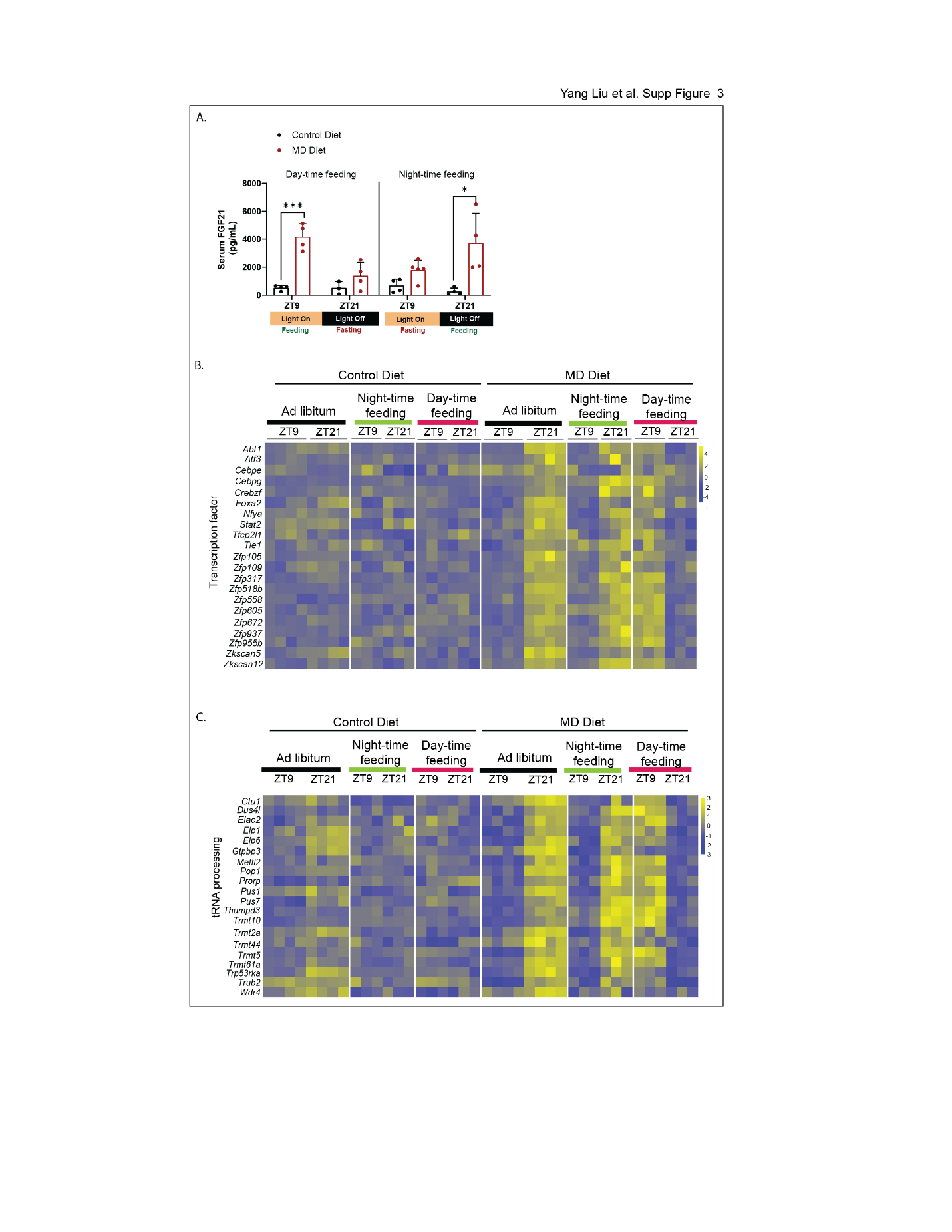

A. Serum levels of FGF21 of six-month old male mice after one week of time restricted feeding on the indicated diet. The column graphs indicate mean value for 3-5 animals at indicated zeitgeber time points. Significance (*p<0.05, ***p<0.001, unpaired t test) is indicated with asterisks.
B. Heatmap depicts the relative expression of transcription factors in animal exposed to time restricted feeding for one week and ad libitum feeding for three weeks.
C. Heatmap depicts the relative expression of genes related to tRNA processing in animal exposed to time restricted feeding for one week and ad libitum feeding for three weeks.

**Figure S4. Rhythmic histone methylation contributes to the reprograming of hepatic circadian rhythms induced by Methionine deprivation, related to Figure 4**

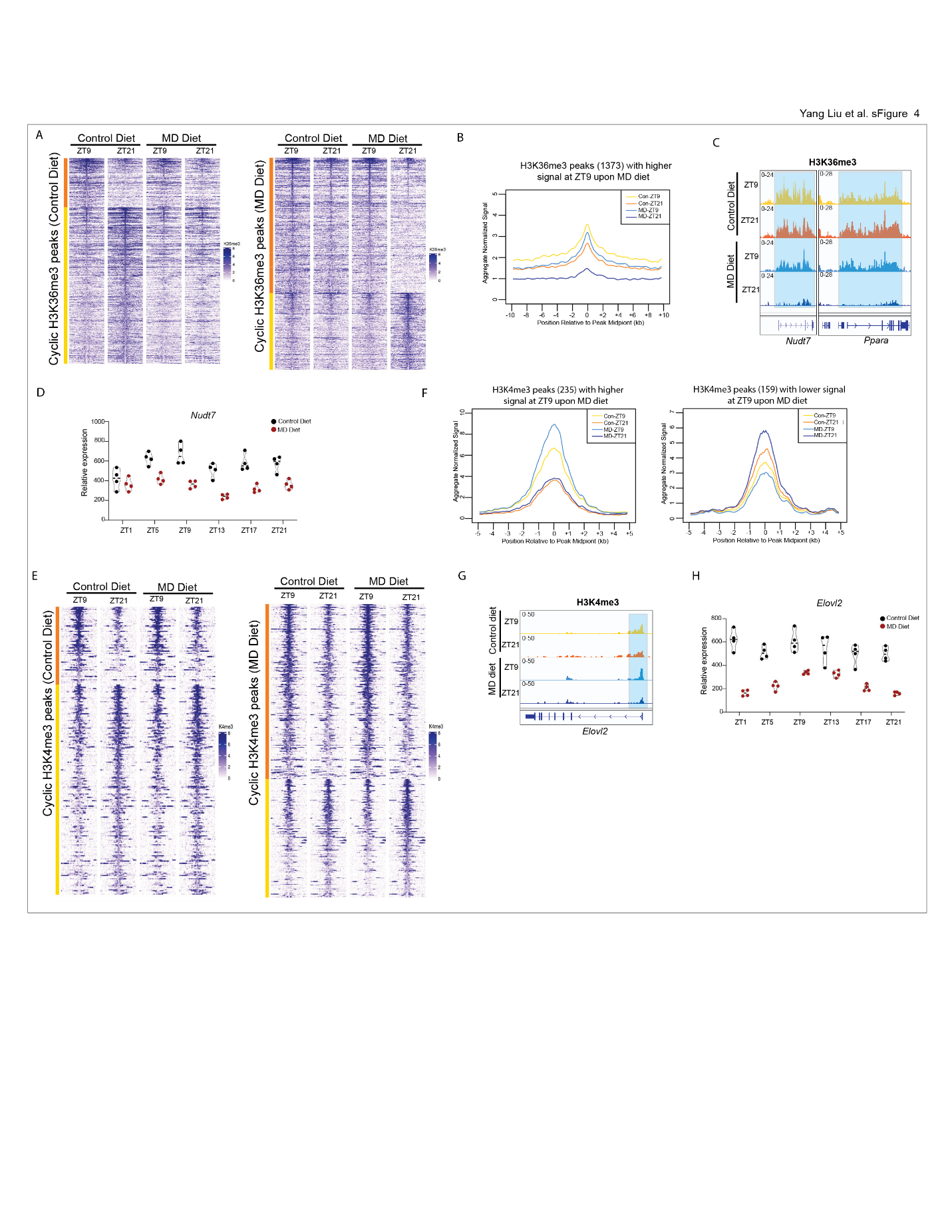

A. The heatmaps depict the H3K36me3 enrichment levels for peaks exhibiting rhythmic oscillation in mice fed on control diet (left panel) and MD diet (right panel). Diet and Zeitgeber time are indicated above the panel, scale bar is to the right. The colored bar to the left of each panel indicates peaks more abundant at ZT9 than ZT21 (orange) or peaks more abundant at ZT21 than ZT9 (yellow).
B. Metagene profile showing H3K36me3 enrichment for 1,373 peaks with a stronger signal at ZT9 compared to ZT21 upon MD diet (peak center ± 10 kb).
C. IGV browser tracks showing H3K36me3 enrichment at *Nudt7* and *Ppara* genes. Diet and Zeitgeber time for animal collection are indicated to the left, gene model is indicated below the data.
D. TMM normalized expression values are presented in the graph for the *Nudt7* gene. Diet group and Zeitgeber time are indicated on the graphs.
E. The heatmaps depict the H3K4me3 enrichment levels peaks exhibiting rhythmic oscillation in mice fed on control diet (left panel) and MD diet (right panel). Diet and Zeitgeber time are indicated above the panel, scale bar is to the right. The colored bar to the left of each panel indicates peaks more abundant at ZT9 than ZT21 (orange) or peaks more abundant at ZT21 than ZT9 (yellow).
F. Metagene profiles showing enrichment for rhythmic H3K4me3 upon MD diet (peak center ± 5 kb).
G. IGV browser tracks showing H3K4me3 enrichment at *Elovl2* genes. Diet and Zeitgeber time for animal collection are indicated to the left, gene model is indicated below the data.
H. TMM normalized expression values are presented in the graph for the *Elovl2* gene. Diet group and Zeitgeber time are indicated on the graphs.

**Figure S5.** **Methionine deprivation reshapes the circadian occupancy of glucocorticoid receptor (GR), related to Figure 5**

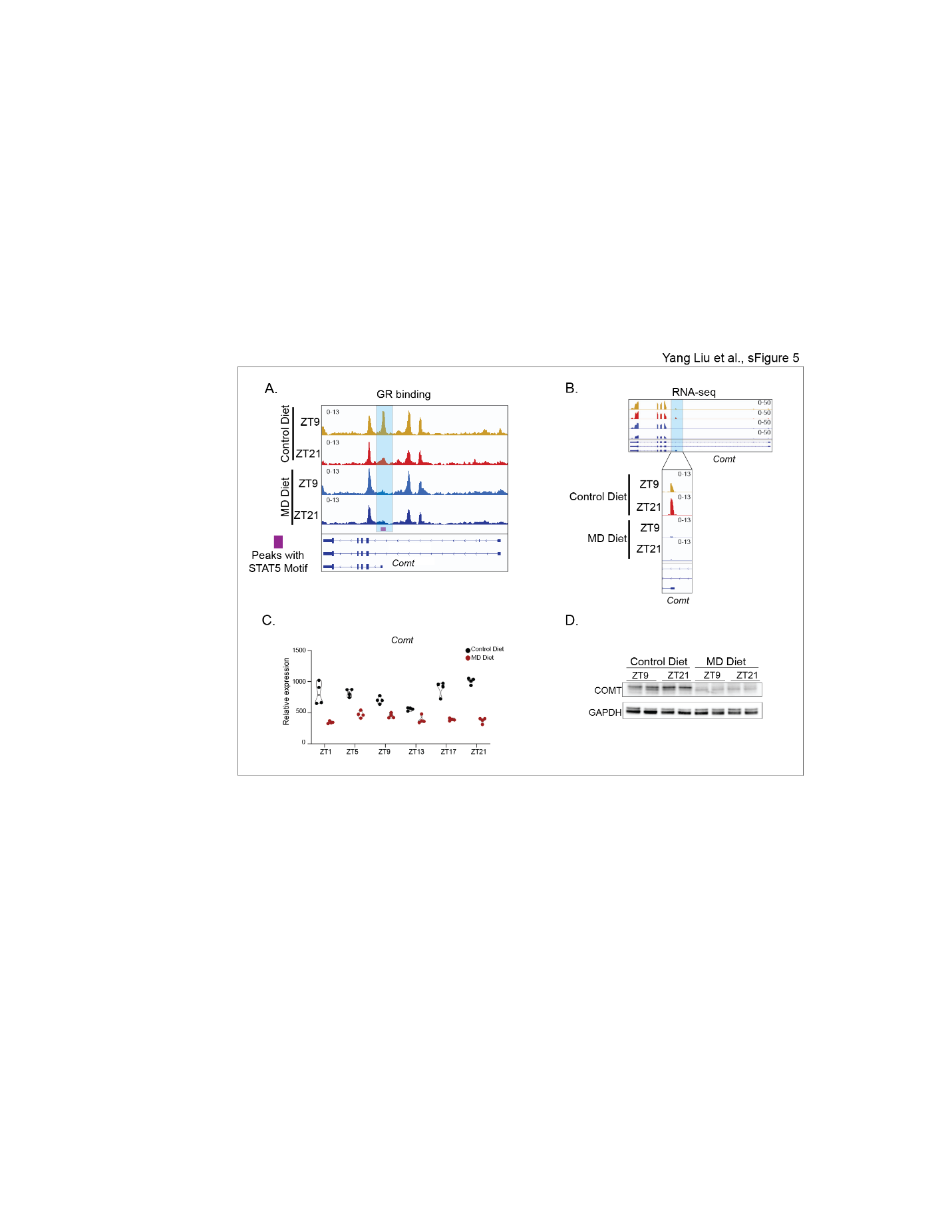

A. IGV browser tracks showing the GR enrichment at *Comt* gene. Diet and Zeitgeber time for animal collection are indicated to the left.
B. IGV browser tracks showing the RNA-seq signal at *Comt* gene.
C. TMM normalized expression values are presented in the graph for the *Comt* gene.
D. The western blot depicts the protein levels of COMT in whole cell lysis of liver from mice on indicated diet for three weeks. GAPDH is included as loading controls

**Figure S6 Altered oscillations of H3K27ac enrichment at GR binding peaks with STAT5 motif, related to Figure 6**

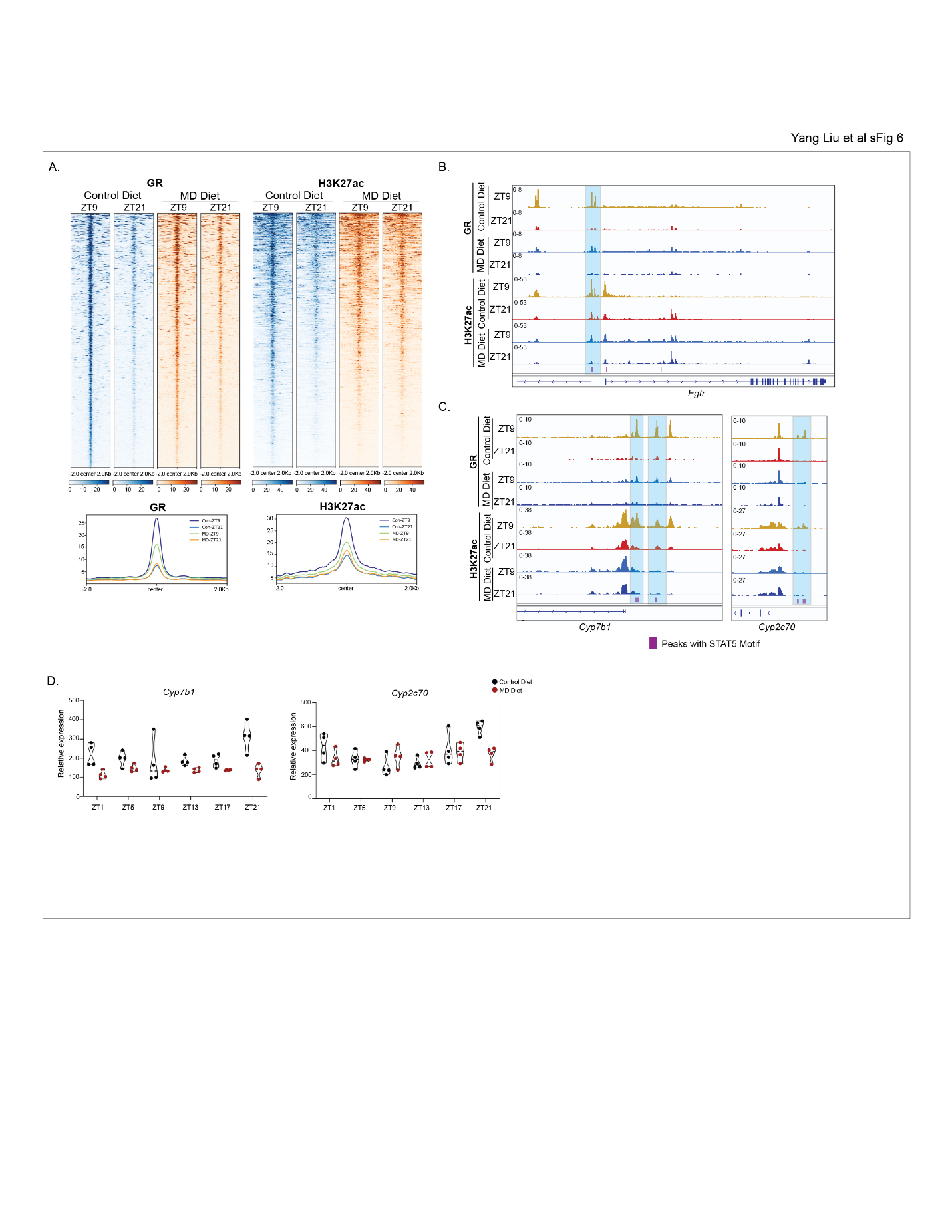

A. The heatmaps and metaplots depict the GR (left) and H3K27ac (right) enrichment levels at 948 rhythmic GR binding peaks with STAT5 motif at ZT9 and ZT21 in mice fed on indicated diet.
B. IGV browser tracks showing the GR and H3K27ac enrichment at *Egfr* genes. Diet and Zeitgeber time for animal collection are indicated to the left.
C. IGV browser tracks showing the GR and H3K27ac enrichment at *Cyp7b1* and *Cyp2c70* genes. Diet and Zeitgeber time for animal collection are indicated to the left.
D. TMM normalized expression values are presented in the graph for the *Cyp7b1* and *Cyp2c70* genes.

**Figure S7 Short-term MD diet changes bile acids levels, related to Figure 6**

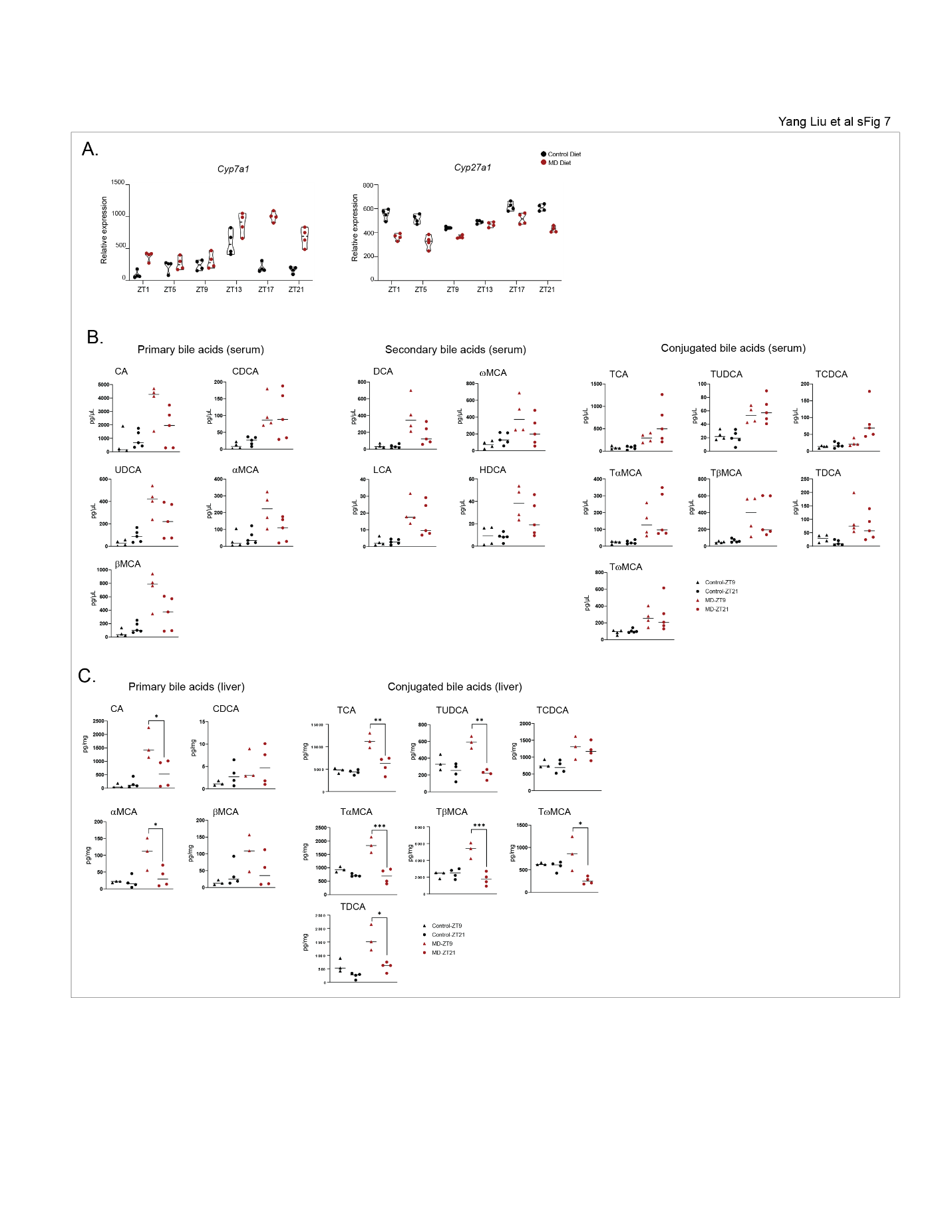

A. TMM normalized expression values are presented in the graph for the *Cyp7a1* and *Cyp27a1* genes.
B. Primary (left), secondary (middle), and conjugated (right) bile acids in serum from mice fed on indicated diet for three weeks.
C. Primary (left) and conjugated (right) bile acids in liver from mice fed on indicated diet for three weeks. Significance (*p<0.05, **p<0.01, ***p<0.001, unpaired t test) is indicated with asterisks.

**Figure S8 Gene ontology analysis for DEGs in liver between GR LKO and control mice, related to Figure 7**

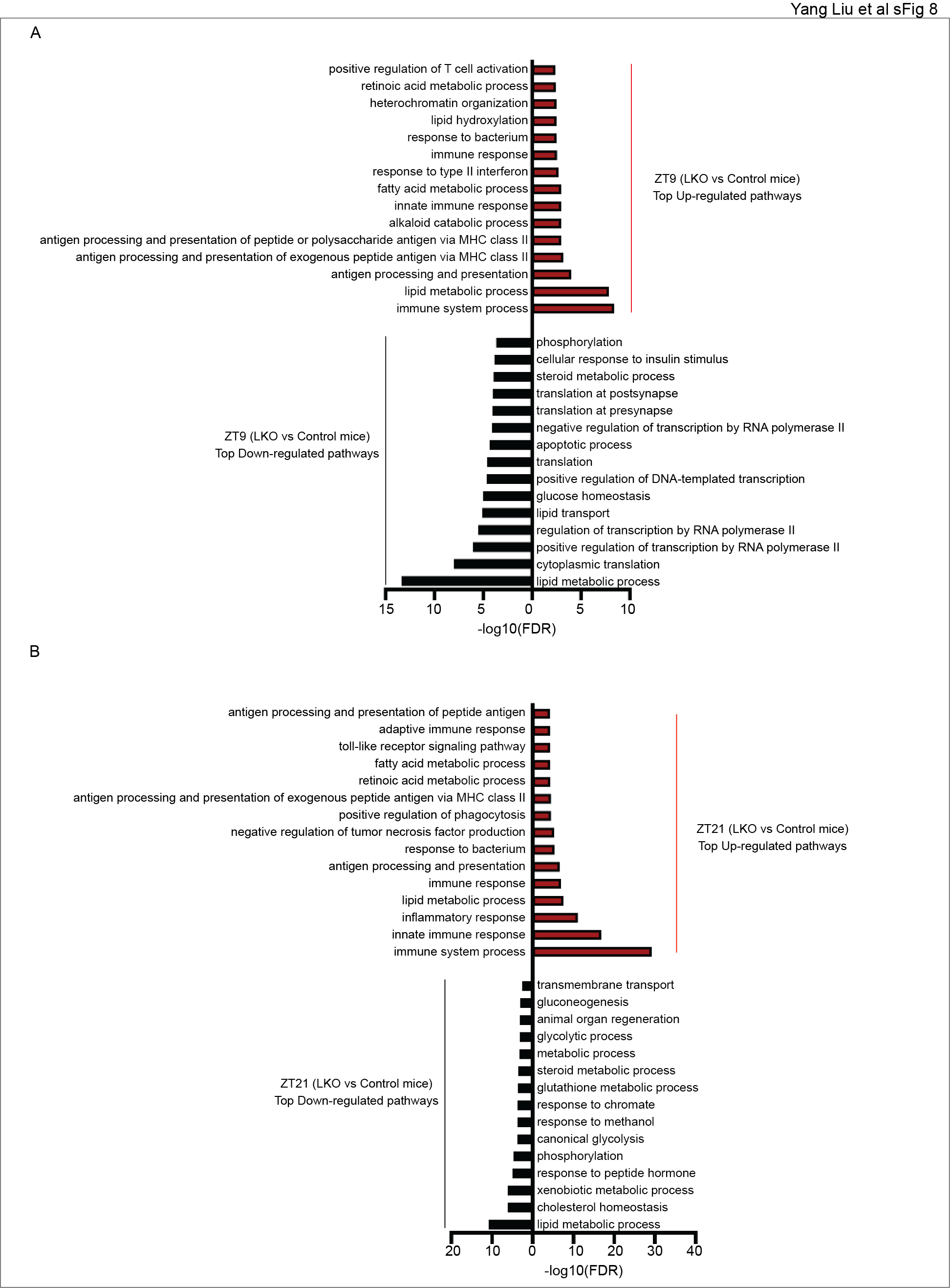

A. Top enriched pathways for DEGs between GR LKO and control mice at ZT9.
B. Top enriched pathways for DEGs between GR LKO and control mice at ZT21.

**Figure S9 GR deletion impairs the circadian rhythms in mice fed on MD diet, related to Figure 7**

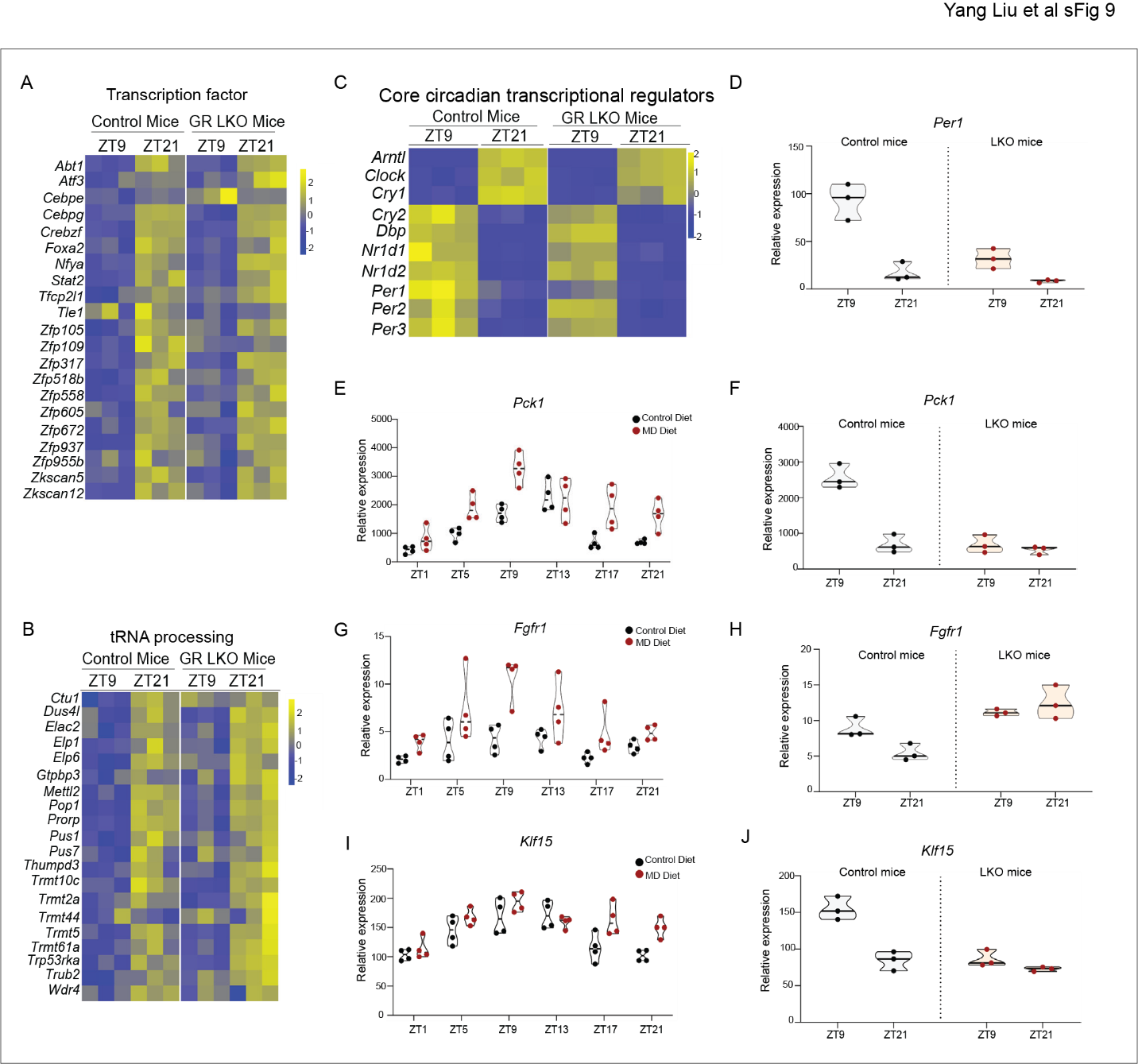

A. Heatmap depicts the relative expression of transcription factors at ZT9 and ZT21 in GR LKO and control mice fed on MD diet for three weeks.
B. Heatmap depicts the relative expression of genes enriched in tRNA processing at ZT9 and ZT21 in GR LKO and control mice fed on MD diet for three weeks.
C. Heatmap depicts the relative expression of core circadian regulators at ZT9 and ZT21 in GR LKO and control mice fed on MD diet for three weeks.
D. TMM normalized expression values of *Per1* gene in liver from GR LKO and control mice on MD diet for three weeks.
E. TMM normalized expression values of *Pck1* gene in liver from wild type mice fed on indicated diet for three weeks.
F. TMM normalized expression values of *Pck1* gene in liver from GR LKO and control mice on MD diet for three weeks.
G. TMM normalized expression values of *Fgfr1* gene in liver from wild type mice fed on indicated diet for three weeks.
H. TMM normalized expression values of *Fgfr1* gene in liver from GR LKO and control mice on MD diet for three weeks.
I. TMM normalized expression values of *Klf15* gene in liver from wild type mice fed on indicated diet for three weeks.
J. TMM normalized expression values of *Klf15* gene in liver from GR LKO and control mice on MD diet for three weeks.
